# *Lnc956*-TRIM28-HSP90B1 complex on replication forks promotes CMG helicase retention to ensure stem cell genomic stability and embryogenesis

**DOI:** 10.1101/2022.03.13.484185

**Authors:** Weidao Zhang, Min Tang, Lin Wang, Hu Zhou, Jing Gao, Zhongliang Chen, Bo Zhao, Ping Zheng

**Author notes:** Correspondence: Ping Zheng, State Key Laboratory of Genetic Resources and Evolution, Kunming Institute of Zoology, Chinese Academy of Sciences, Kunming, Yunnan 650223, China., Bo Zhao, Kunming Institute of Zoology, Chinese Academy of Sciences, Kunming, Yunnan 650223, China. These authors contributed equally (Weidao Zhang, Min Tang).

## Abstract

Replication stress is a major source of endogenous DNA damage. Despite that numerous proteins have been identified on replication forks to modulate fork or replication machinery activities, it remains unexplored whether non-coding RNAs can localize on stalled forks and play critical regulatory roles. Here we identify an uncharacterized lncRNA NONMMUT028956 (*Lnc956* for short) predominantly expressed in mouse embryonic stem cells. *Lnc956* is recruited to stalled replication forks to prevent fork collapse and preserve genomic stability, and is essential for mouse embryogenesis. Mechanistically, it drives assembly of the *Lnc956*-TRIM28-HSP90B1 ribonucleoprotein (RNP) complex on stalled forks in an inter-dependent manner downstream of ATR signaling. This RNP complex physically associates with MCM2-7 hexamer via TRIM28 and directly regulates the CMG helicase retention on chromatin. The regulation of RNP on CMG retention is mediated by HSP90B1’s chaperoning function. These findings reveal a novel pathway which actively regulates replisome retention to prevent fork collapse.

## Introduction

Pluripotent stem cells (PSCs) are capable of self-renewal and differentiation into all cell types in the body, and are cellular basis for organism development. Due to their unique functions, maintaining stable genome is fundamental for stem cells. Perturbations on genome stability can cause stem cell apoptosis, impair differentiation potentials, and induce the tumorigenicity ^1, 2, 3^. It has been well recognized that PSCs possess superior stable genome than differentiated cells. For instance, mouse embryonic stem cells (ESCs) display 100-fold lower mutation rate than their isogenic mouse embryonic fibroblasts (MEFs) ^4^. How this is achieved in PSCs remains largely unknown. Understanding the mechanisms can not only provide valuable insights into the developmental failure/defects, but also help address the issue of genomic instability seen in cultured PSCs, which hampers the full applications of PSCs in cell-based regenerative medicine.

Limited studies suggested that PSCs employ unique strategies and regulators to efficiently safeguard the genomic stability in a range of cellular activities. For example, a homologous recombination (HR)-based way which involves PSC-specific proteins ZSCAN4 and DCAF11 operates in telomere lengthening ^5, 6^. In addition, telomere protection in ESCs does not require TRF2, which is instead essential in somatic cells ^7, 8^. In response to DNA double strand breaks (DSBs), PSCs prefer to choose HR-mediated repair pathway ^9^. What determine this pathway choice remains mysterious, but PSC-specific proteins SALL4 and Filia were identified to play parts in HR repair processes ^10, 11^. DNA replication stress represents a major source of endogenous DNA damages and imposes big threat to genomic integrity ^12^. PSCs contain high level of DNA replication stress due to the lack of G1 checkpoint ^13^, but they are able to efficiently resolve replication stress ^14^. Although our recent study identified a PSC-specific Filia-Floped protein complex on replication forks as a regulator ^14^, the underlying mechanisms are complicative and far from clear.

DNA replication is carried out by a large molecular machine called replisome. The core components of replisome include the CDC45-MCM-GINS (CMG) DNA helicase, PCNA, and replicative DNA polymerases. In addition, numerous auxiliary factors associate with replisome to modulate its assembly, disassembly, and activities under unperturbed and perturbed conditions in a context-dependent manner ^15^. CMG helicase, which is composed of CDC45, minichromosome maintenance proteins 2 to 7 (MCM2-7), and the go-ichi-ni-san (GINS) complex (comprising SLD5, PSF1, PSF2 and PSF3), plays rate limiting role in DNA replication by unwinding DNA along the leading strand template. Upon replication fork stalling, the intra-S phase replication checkpoint mediated by kinase ataxia telangiectasia and Rad3-related (ATR) is activated, and the local and global responses coordinate to preserve stalled forks and promote fork restart after obstacles are removed. Failure to repair and restore the stalling forks will cause fork collapse, which is defined as the inability to restart forks and characterized by the breakage of DNA into DSBs as well as the dissociation of replisomes from forks ^16, 17, 18^. Thanks to the innovative methods such as isolation proteins on nascent DNA (iPOND) and nascent chromatin capture (NCC) ^19, 20^, which coupled with mass spectrometry have provided systematic insights into the dynamic protein compositions and regulations of replication forks under normal and stressed conditions ^17, 21, 22^. While there is general consensus that the replication checkpoint regulates fork repair, whether checkpoint can also directly stabilize replisome to prevent fork collapse remains uncertain. Several studies in vertebrates proposed a checkpoint-dependent replisome stability model ^23, 24^. However, a recent study using improved iPOND protocol combined with SILAC mass spectrometry suggested that the major functions of ATR replication checkpoint on stalled forks are to repair forks rather than to stabilize replisomes in mammalian cells. The replisome core component CMG helicase remained stable on forks until the late time of stress treatment even if the checkpoint was inactivated ^25^. This observation implied that replisome dissociation from stalled forks could simply be the consequence of fork collapse. Thus, whether replisome stability underlies the replication fork collapse still remains controversial.

To date, only protein regulators have been reported on replication forks. It is unclear if non-coding RNAs can reside on replication forks and regulate forks or replisomes. Long non-coding RNAs (lncRNAs) are able to bind to DNA, RNAs, or proteins, can drive highly efficient condensation of interacting proteins and largely enhance the protein functions ^26, 27^. In light of the knowledge, we wondered whether lncRNAs are associated with replication forks in PSCs and lncRNA-mediated regulation attributes to the efficient replication stress response in PSCs. In this study, we modified iPOND method in order to isolate fork-associated RNA species and named this method as isolate <u>R</u>NAs <u>o</u>n <u>n</u>ascent <u>D</u>NA (iROND). Using iROND coupled with RNA-seq, we systematically analyzed the lncRNA species localized on replication forks of differentiated cells and PSCs under normal and stressful conditions. Further, we characterized in detail one ESC-prevalent lncRNA *Lnc956* (NONCODE ID: NONMMUT028956, *Lnc956* for short). Our results showed that ATR replication checkpoint induced *Lnc956*-TRIM28-HSP90B1 ribonucleoprotein (RNP) complex assembly on stalled forks under replication stress. This RNP complex promotes CMG helicase retention on chromatin to prevent replication fork collapse. Thus, our study suggested that ATR replication checkpoint might regulate the retention of replisome on chromatin, which underlies fork collapse in certain cellular context, for instance, in stem cells.

## Results

### ESC-prevalent lncRNA *Lnc956* resides on replication forks and is essential for fork restart after replication stress

To find out if there are lncRNAs on replication forks, we modified the iPOND protocol by avoiding RNase contamination and named this method as iROND (isolate <u>R</u>NAs <u>o</u>n <u>n</u>ascent <u>D</u>NA) (Supplementary Fig. 1a). iROND samples were prepared from mouse ESCs and NIH3T3 cells under unperturbed or hydroxylurea (HU) treatment condition in two replicates, and were subject to RNA extraction followed by RNA sequencing. LncRNAs were annotated according to the NONCODE database (http://www.noncode.org/). Interestingly, three uncharacterized lncRNAs (NONMMUT060689, NONMMUT028956, NONMMUT006458) were reproducibly identified on replication forks of ESCs, whereas no lncRNA was repeatedly detected in iROND samples of NIH3T3 (Fig. 1a, Supplementary data 1). Quantitative RT-PCR (qRT-PCR) showed that among the three lncRNAs, NONMMUT060689 and NONMMUT028956 were prevalently expressed in ESCs when compared to differentiated cells (Supplementary Fig. 1b) or various tissue samples (Supplementary Fig. 1c), and the level in ESCs did not display drastic induction after HU treatment (Supplementary Fig. 1b). These observations suggested that stable association of some lncRNAs with replication forks might be unique in ESCs. We then took the ESC-specific lncRNA NONMMUT028956 (*Lnc956* for short) as an example to understand this phenomenon in detail.

**Fig 1.**
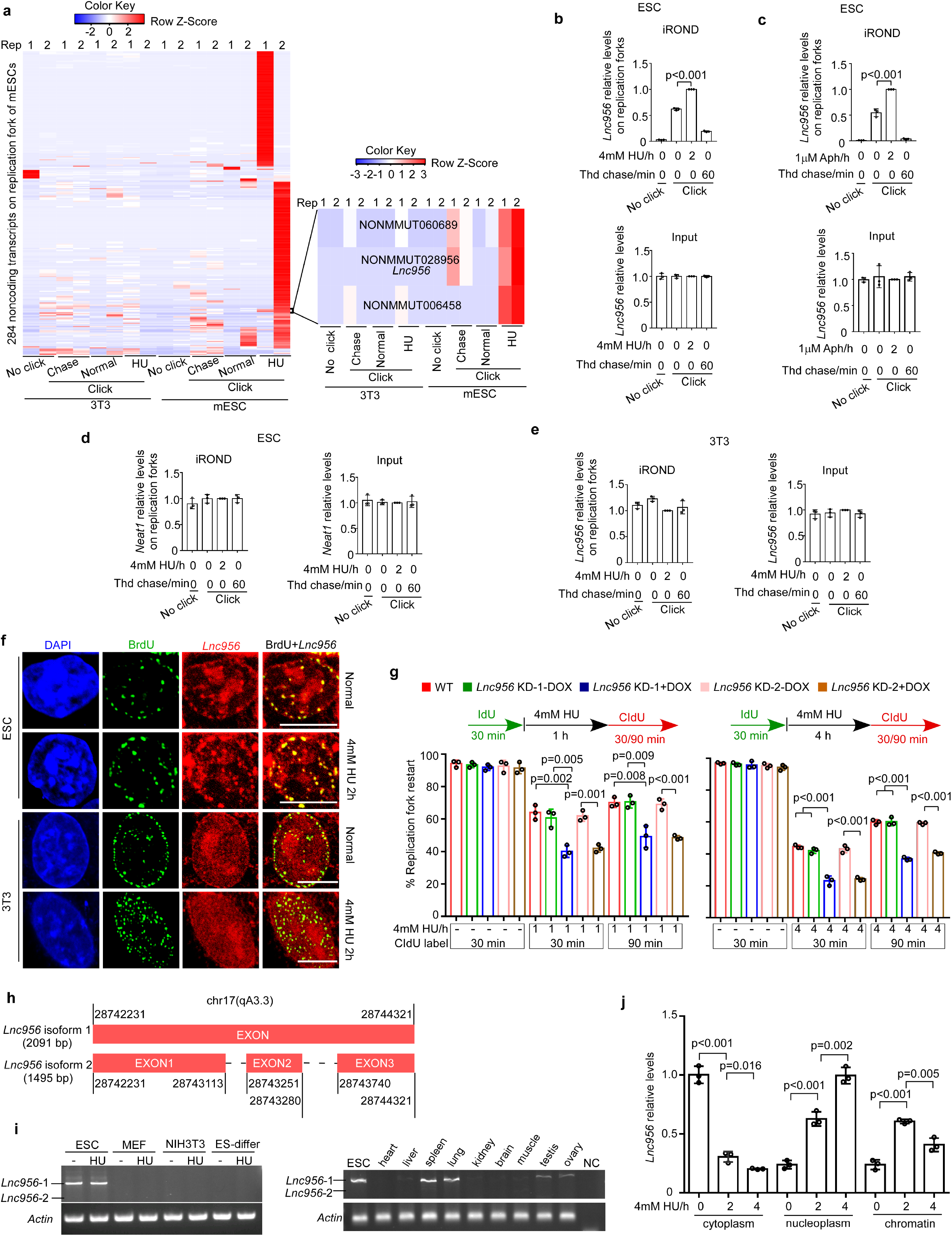
*Lnc956* resides on replication forks of mouse ESCs and promotes stalled fork restart. **a** Two replicative iROND samples were prepared from NIH3T3 cells and mouse ESCs (mESCs) under different experimental conditions. RNA-seq analyses of iROND samples reproducibly detected several lncRNAs on isolated replication forks of ESCs under hydroxyurea (HU) treatment condition. All cells were cultured under normal condition (normal) or treated with 4 mM HU for 2 h. Samples without click reaction (no click) or with 10 μM thymidine chase (Thd) were included. Rep represents repeat. **b-c** Replication forks were isolated and the relative levels of *Lnc956* resided on forks were determined by quantitative RT-PCR (qRT-PCR). *Lnc956* were detected on replication forks and the levels were increased by HU (**b**) or aphidicolin (Aph) (**c**) treatment. The levels of *Lnc956* in inputs were normalized by *Actb*. **d** The relative levels of lncRNA *Neat1* resided on replication forks of mESCs were determined as in b. *Neat1* was not detected in replication forks. **e** The relative levels of *Lnc956* resided on replication forks of NIH3T3 cells were determined as in b. No *Lnc956* was detected. In b-e, data were from three independent biological repeats. **f** Nascent DNA was labeled by BrdU for 5 min. RNA-FISH revealed the co-localization of the *Lnc956* with nascent DNA under normal and HU treatment conditions, indicating the allocation of *Lnc956* on replication forks in ESCs. However, there was no obvious co-localization in NIH3T3 cell. Scale bars, 10 μm. Three independent experiments were repeated with similar results. **g** ESCs were treated with HU for 1 h (left panel) or 4 h (right panel). DNA fiber assay revealed that doxycycline (DOX)-induced knock down (KD) of *Lnc956* impaired the stalled fork restart at both conditions. At least 200 fibers from three independent experiments were analyzed. **h** The chromosomal location information of the two isforms of *Lnc956*. Red boxes indicate exons. **i** RT-PCR showed the prevalent expression of long isoform of *Lnc956* in ESCs and other differentiated cells (left panel) or tissues (right panel). ES-differ refers to the ESC-differentiated cells in the presence of retinoid acid. Three independent experiments were repeated with similar results. **j** *Lnc956* displayed dynamic redistribution in compartments of cytoplasm, nucleoplasm and chromatin after HU treatment. All data were shown as mean ± SEM from three replications, two-tailed Student’s *t*-test.

We first validated the localization of *Lnc956* on replication forks by qRT-PCR analysis of iROND precipitates. As shown, *Lnc956* was enriched on replication forks compared to non-click and one-hour thymidine chase negative controls. Moreover, replication stress induced by HU (Fig. 1b) or aphidicolin (Fig. 1c) treatment increased the *Lnc956* level on forks. As a control, lncRNA *Neat1*, which localizes on nuclear paraspeckles ^28^, was not detected on replication forks of ESCs (Fig. 1d). In sharp contrast to the observation on ESCs, *Lnc956* was not detected on replication forks of NIH3T3 cells (Fig. 1e). We performed RNA fluorescence in situ hybridization (FISH) combined with BrdU immunostaining to validate the subcellular localization. In mESCs, almost all nascent DNA fragments pulse-labeled with BrdU were co-localized with *Lnc956* foci under the normal and HU treatment conditions. However, few *Lnc956* foci showed co-localization with BrdU foci in NIH3T3 cells (Fig. 1f). We next investigated if *Lnc956* has functions on replication forks. To this end, we efficiently knocked it down with two independent doxycycline (DOX)-inducible short hairpin RNAs (shRNAs) (Supplementary Fig. 1d), and examined the replication fork behaviors under normal and HU treatment conditions using DNA fiber assay ^29^. Under normal culture condition, *Lnc956* knock down (KD) had no visible influence on replication fork speed (Supplementary Fig. 1e) or fork symmetry (Supplementary Fig. 1f). Concordantly, cell cycle in *Lnc956* KD ESCs was not altered (Supplementary Fig. 1g). However, after HU treatment for 1 hour (h) or 4 hours, *Lnc956* KD severely impaired the stalled fork restart (Fig. 1g). Similar result was obtained when replication stress was evoked by aphidicolin treatment (Supplementary Fig. 1h). In line with the fork restart impairment, *Lnc956* KD enhanced the fork asymmetry under replication stress condition (Supplementary Fig. 1i). Taken together, these results suggested that fork-allocated *Lnc956* plays essential role in promoting DNA replication under stressed conditions.

To better understand *Lnc956*, we obtained its full length by 5’-end and 3’-end RACE (rapid amplification of cDNA ends). Intriguingly, two isoforms were identified (Fig. 1h), with the longer isoform being dominant in both ESCs and tissue samples (Fig. 1i). To examine if *Lnc956* undergo subcellular translocation upon replication stress, we fractionated the cellular compartments of ESCs into cytoplasm, soluble nucleoplasm and insoluble chromatin. qRT-PCR analysis revealed a dynamic relocation from cytoplasm to nucleus (Fig. 1j). We also examined the coding ability of several potential coding frames in *Lnc956* and failed to detect any translation of protein or peptide (Supplementary Fig. 1j). In addition, *Lnc956* is predicted to be partially conserved among mammalian species (Supplementary Fig. 1k). Taken together, we identified a functional lncRNA which shows preferential expression in ESCs and resides on replication forks to facilitate stalled fork restart and promote DNA replication under stressful conditions.

### *Lnc956* promotes the retention of CMG helicase on chromatin to maintain genomic stability under replication stress

Depletion of *Lnc956* compromised the stalled fork restart following replication stress, indicating that stalled forks were prone to collapse in the absence of *Lnc956*. Fork collapse is associated with replisome dissociation and DNA DSB formation. We went on to examined the influences of *Lnc956* depletion on dynamics of the two events after HU treatment. We purified nascent DNA by iPOND and evaluated the amounts of replisome core components including CMG components and PCNA on forks. The levels of these proteins were comparable between *Lnc956* proficient and deficient ESCs under unperturbed condition. Upon HU treatment, the CMG components and PCNA in *Lnc956* proficient ESCs remained stable on replication forks at initial 1 h and 2 h of HU treatment, but displayed obvious dissociation at 4 h of treatment. Dissociation of these proteins after 4 h HU treatment was not due to the completion of DNA replication (Supplementary Fig. 2a). Rather, it might reflect the outcome of fork breakage. Compared to *Lnc956* proficient ESCs, the dissociation of CMG components was drastically accelerated in *Lnc956* KD cells. At 1 h of HU treatment, CMG components already displayed obvious unloading in both KD cells. PCNA also dissociated at 2 h of HU treatment (Fig. 2a, Supplementary Fig. 2b). We further isolated chromatin and re-evaluated the chromatin binding of CMG components and PCNA. Consistent dynamics was detected (Fig. 2b, Supplementary Fig. 2c). Moreover, recovery for 30 mins and 90 mins following HU treatment (1 h or 4 h) as depicted in Fig. 1g did not change the CMG dissociation pattern (Supplementary Fig. 2d, e). We then examined the dynamics of DNA DSB formation indicative of fork breakage under the same treatment conditions. Compared to *Lnc956* proficient ESCs*, Lnc956* KD ESCs contained higher basal level of DNA DSBs under normal culture condition. HU treatment induced DSB formation in all ESC groups. However, *Lnc956* KD had no influence on the kinetics of fork breakage and DSB formation under stressful condition. Only at 4 h of HU treatment, were the stalled forks broken down to form DSBs in a manner irrelevant to *Lnc956* KD (Fig. 2c). Thus, the stress-induced replisome unloading coincided with the DSBs formation in WT ESCs, whereas replisome dissociation occurred before fork breakage in *Lnc956* KD ESCs. These results suggested that replisome dissociation and fork breakage were separately regulated, and *Lnc956* might directly regulate the CMG (or replisome) retention on replication forks under stressful conditions.

**Fig 2.**
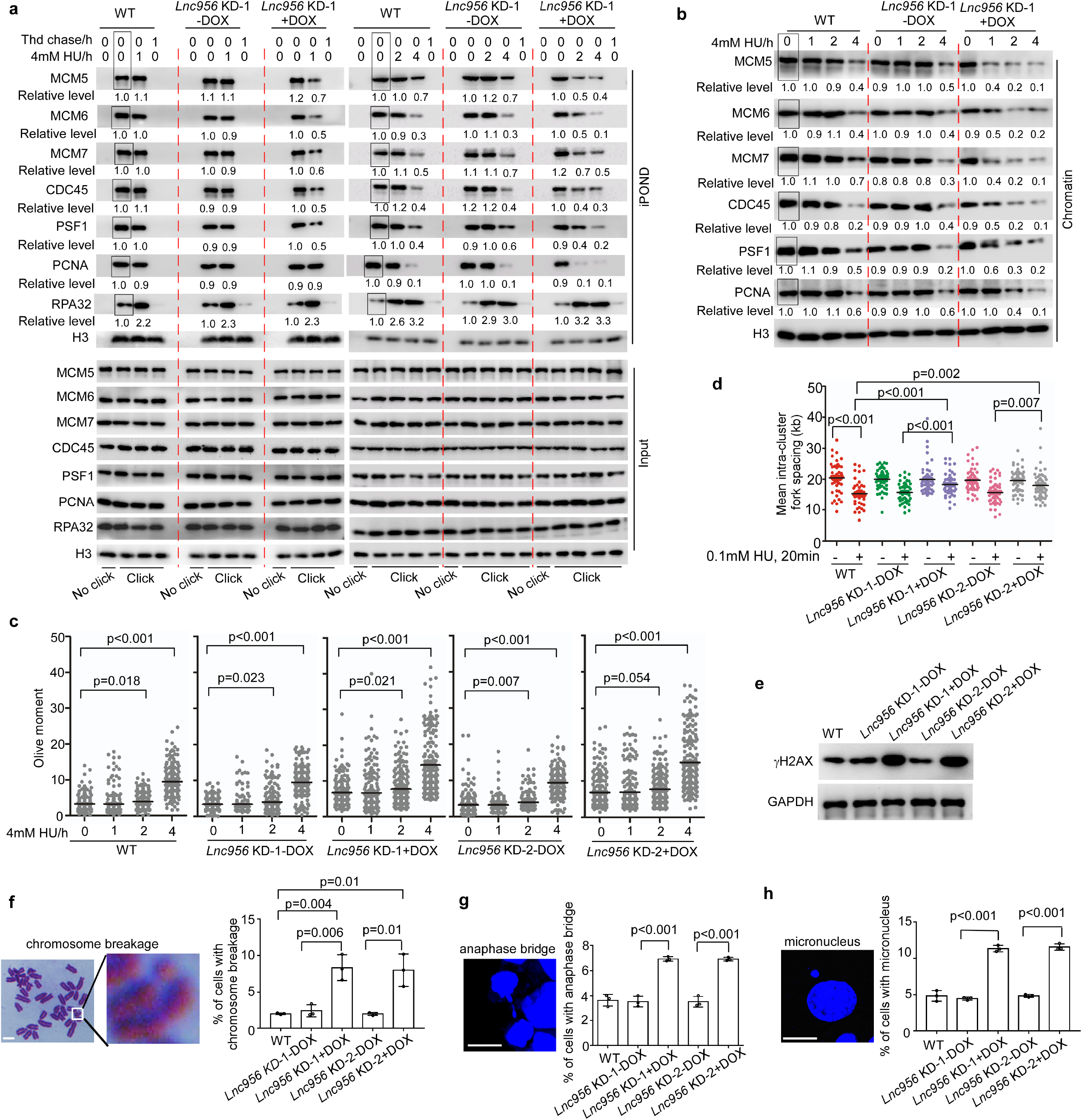
*Lnc956* promotes the CMG helicase retention on chromatin to ensure genomic stability under replication stress. **a** *Lnc956* KD (KD-1) accelerated unloading of CMG components (MCM5, 6, 7, CDC45 and PSF1) from replication forks under HU treatment condition. The premature unloading appeared at as early as 1 h following HU treatment. Other replication fork proteins PCNA and RPA32 displayed different kinetics. Samples without click reaction (no click) and with 10 μM thymidine (Thd) chase were included as negative controls for iPOND. **b** Consistently, *Lnc956* KD (KD-1) accelerated the dissociation of CMG components from chromatin. In (a, b), the relative protein levels were normalized by histone H3, and the levels in samples marked with box were set as 1. All experiments were repeated three times with similar results. **c** Although *Lnc956* KD accelerated the unloading of CMG, it did not affect the dynamics of HU-induced DNA DSB formation measured by neutral comet assay. At least 200 tails were analyzed in each group. Data were from three independent replications. **d** *Lnc956* KD reduced dormant origin firing as indicated by the increased inter-origin distances. At least 50 continuous fibers from three independent replications were analyzed. **e** Compared to WT ESCs, *Lnc956* KD cells accumulated more DNA damages as indicated by higher level of γH2AX. The experiments were repeated three times with similar results. **f** *Lnc956* KD mESCs had higher rate of chromosome breakage. Representative image of chromosome breakage was shown in Left panel and the quantification (right panel) was obtained from at least 50 metaphase spreads in three independent replications. **g** *Lnc956* KD increased the rate of anaphase bridge. The representative image (left panel) and quantification (right panel) of anaphase bridge were shown. At least 50 anaphase-stage cells from three independent replications were analyzed in each group. **h** *Lnc956* KD stimulated micronuclei formation. The representative image (left panel) and quantification (right panel) of micronucleus were shown. At least 50 visual fields containing 1000 cells were analyzed in three replications in each group. All data were shown as mean ± SEM, two-tailed Student’s *t*-test. Scale bar, 10 μm.

We went on to gain more insights into the influences of *Lnc956* depletion on replication checkpoint. The ATR-CHK1 signaling central to checkpoint was normally activated in *Lnc956* KD ESCs (Supplementary Fig. 2f). No nascent DNA degradation was observed, indicating that *Lnc956* was not involved in fork protection (Supplementary Fig. 2g). However, dormant origin firing was compromised by *Lnc956* KD, as indicated by the increases in inter-origin distances (Fig. 2d). This observation implicated that the dormant origin density might be reduced upon replication stress owing to the dissociation of MCM hexamers in the absence of *Lnc956*.

CMG helicase plays rate-limiting role in DNA replication. Its core component MCM hexamer is loaded on chromatin in G1 phase and can not be reloaded during S phase. Undesirable dissociation of MCM from chromatin during S phase disables DNA replication restart leading to fork collapse. In addition, it reduces dormant origin firing and hampers the completion of DNA replication, which can be manifested by chromosomal breakages, chromosomal bridges during mitosis, and micronuclei formation ^30^. Indeed, *Lnc956* KD ESCs contained more DNA DSBs under normal culture conditions, as measured by immunoblotting analysis of γH2AX (Fig. 2e). The frequencies of chromosomal breakages (Fig. 2f), mitotic chromosomal bridges (Fig. 2g), and micronuclei formation (Fig. 2h) were significantly elevated in *Lnc956* KD ESCs.

### Loss of *Lnc956* causes female-biased embryo lethality

DNA replication-associated genomic instability often affects embryogenesis. For instance, *MCM4^chaos^*^3^ mutation distabilizes MCM2-7 complex and significantly reduces the chromatin-associated hexamers, resulting in genomic instability and embryonic lethality ^31, 32^. Based on the genomic instability phenotypes observed in ESCs, we speculated that *Lnc956* might play essential roles in embryo development. To test this hypothesis, we generated *Lnc956* knockout (KO) mice by CRISPR/Cas9 mediated mutation strategy (Supplementary Fig. 3a). Two KO mouse lines were obtained from the same targeting design. *Lnc956* KO was validated by Sanger sequencing of amplified DNA fragment (Supplementary Fig. 3b), and genotyped by PCR examination of tail DNA samples (Supplementary Fig. 3c). Because the two mouse lines harbored similar fragment deletion (Supplementary Fig. 3b), we utilized line 1 for detailed investigations. No gross abnormality was detected in heterozygous mice. However, crossings of heterozygous mice did not produce offsprings with normal Mendelian ratio, in which the birth of homozygotes was severely compromised (Fig. 3a). Other breeding strategies (*Lnc956*^-/-^ females x *Lnc956*^+/-^ males, and *Lnc956*^+/-^ females × *Lnc956*^-/-^ males) led to the consistent results (Fig. 3b). Thus, complete loss of *Lnc956* caused embryonic lethality. Intrigingly, embryo death displayed obvious gender bias. KO females were markedly under-represented compared with KO males. Based on the Mendelian ratio, we estimated that *Lnc956* KO caused lethality in around 50% of male embryos, and in about 85% of female embryos, respectively, in all three mating sets (Fig. 3a, b). It should be noticed that heterozygous offsprings, of which either one parental genotype was homozygous null (*Lnc956*^+/-^ females × *Lnc956*^-/-^ males, or *Lnc956*^-/-^ females × *Lnc956*^+/-^ males), also displayed female-biased under-representation (Fig. 3b). This observation suggested that heterozygous loss of *Lnc956* might also lead to female embryo death.

**Fig. 3.**
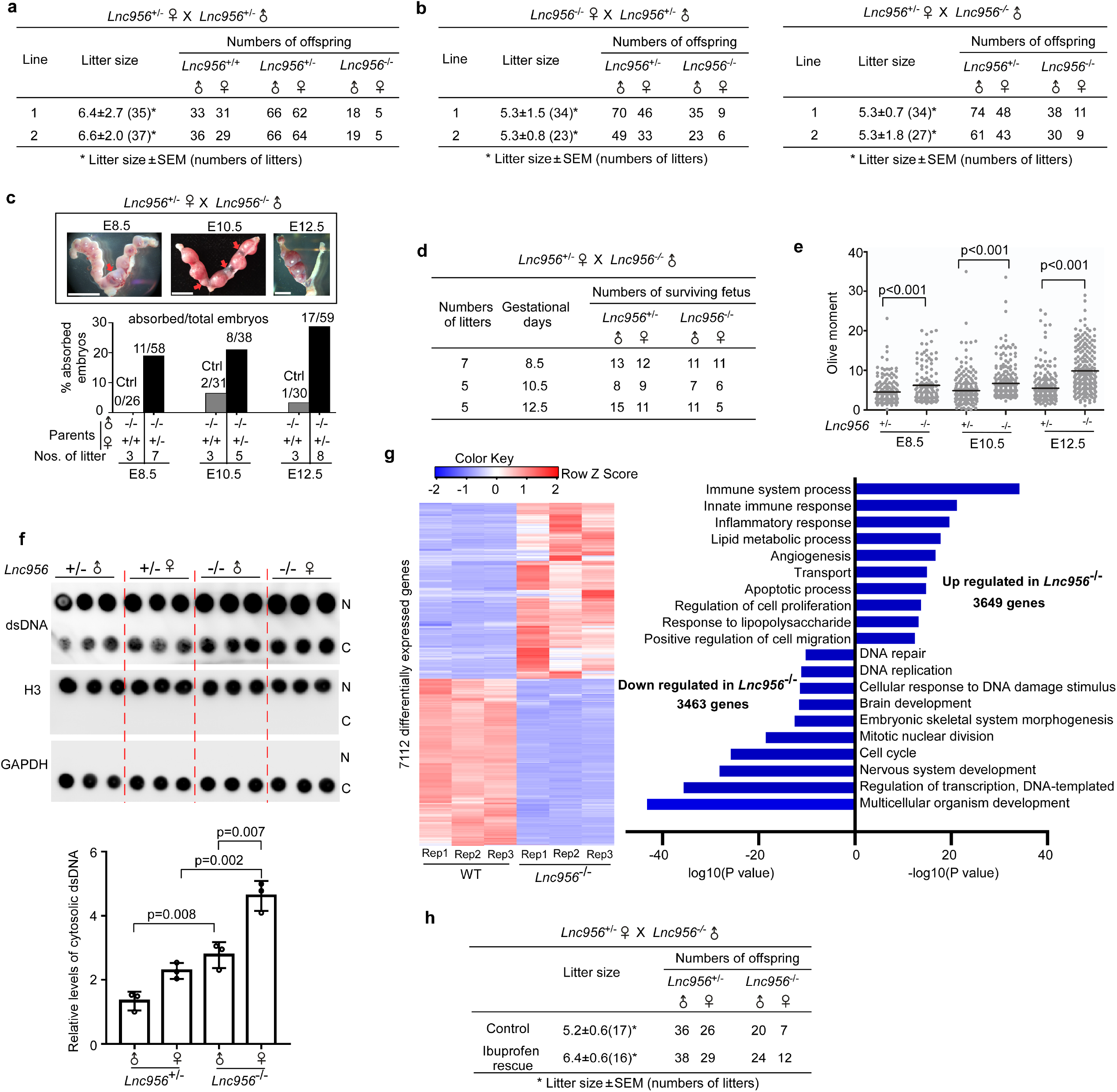
*Lnc956* knockout (KO) causes female-biased embryo lethality. **a-b** Mating experiments with different strategies revealed that complete loss of *Lnc956* caused female-biased embryonic death. **c** Representative images of uterus collected from *Lnc956*^+/-^ pregnant females mated with *Lnc956*^-/-^ males at embryonic day 8.5 (E8.5), E10.5 and E12.5 (upper panel). Red arrows indicated the absorbed dead embryos. Lower panel showed the percentages of absorbed embryos at each developmental stage. The mating (female *Lnc956*^+/+^ × male *Lnc956*^-/-^) was used as control (Ctrl). Scale bars, 1 cm. **d** Genotyping of the survived embryos showed that female was under-represented in homozygous null at E12.5. **e** Single cells dissociated from embryos at different embryonic stages were subject to neutral comet assay. Cells from *Lnc956*^-/-^ embryos contained higher level of DNA DSBs than those from *Lnc956*^+/-^ counterparts. At least 200 tails were analyzed in each group. Data were from three independent replications. **f** Cells were dissociated from E12.5 embryos collected from *Lnc956*^+/-^ pregnant females mated with *Lnc956*^-/-^ males. Dot immunoblotting (upper panel) showed that *Lnc956*^-/-^ embryonic cells contained higher level of cytosolic double strand DNA (dsDNA) than *Lnc956*^+/-^ cells. Lower panel showed the relative levels of cytosolic dsDNA from three independent replications. N refers to nucleus, and C refers to cytoplasm. **g** Heatmap (left panel) and Gene ontology (GO) enrichments (right panel) of differentially expressed genes (DEGs) between wildtype and *Lnc956*^-/-^ embryos at E12.5. **h** Administration of ibuprofen (0.4 mg/mL in drinking water) to pregnant females from gestation day 7.5 onward could partically rescue the female-biased embryonic death of KO mice. All data were shown as mean ± SEM, two-tailed Student’s *t*-test.

We next examined the time window when the embryos die. *Lnc956*^+/-^ females were mated with *Lnc956*^-/-^ males and the blastocysts were collected from uterus. The genotypes of blastocysts display normal Mendelian ratio, indicating that *Lnc956* KO did not impair the pre-implantation development. We then collected post-implantation embryos from embryonic day 8.5 (E8.5) to E12.5. Compared to the control (*Lnc956*^+/+^ females × *Lnc956*^-/-^ males), the increase in embryo absorption was observed at all stages in the mating group of *Lnc956*^+/-^ females × *Lnc956*^-/-^ males (Fig. 3c). This observation implicated that embryo death may occur at variable stages of post-implantation. We further analyzed the genotype as well as the gender of the survived embryos. In line with the mating data, *Lnc956*^-/-^ embryos were under-represented at three developmental stages and the tendency increased with the advanced gestation (Fig. 3d). Of interest, the obvious gender bias was detected only at E12.5 when the sex determination has occurred (Fig. 3d).

The KO embryo contained elevated DNA DSBs than the *Lnc956*^+/-^ counterparts (Fig. 3e). DNA replication-associated genomic instability is usually characterized by the formation of cytosolic DNA fragments, which are able to activate the cGAS-STING pathway and lethal inflammation ^32^. We then examined whether *Lnc956* KO increased the level of cytosolic micronuclei and induced inflammation response in embryos. E12.5 embryos were collected from *Lnc956*^+/-^ pregnant females mated with *Lnc956*^-/-^ males. Dot immunoblotting showed that cells from *Lnc956*^-/-^ embryos contained higher level of cytosolic double strand DNA (dsDNA) than those from *Lnc956*^+/-^ counterparts (Fig. 3f). Further we performed RNA-seq to compare the mRNA expressions between WT and *Lnc956*^-/-^ embryos at E12.5. Total 7112 differentially expressed genes (DEGs) were identified to show consistent up- or down-regulation between two groups (fold change ≥2) (Fig. 3g, Supplementary data 2). Gene ontology (GO) enrichment analyses of these DEGs revealed that innate immune response and inflammation response were the top processes enriched in DEGs up-regulated in *Lnc956*^-/-^ embryos, whereas processes including cellular response to DNA damage stimulus, DNA replication and repair were enriched in DEGs down-regulated in *Lnc956*^-/-^ embryos (Fig. 3g, Supplementary Fig. 4). These lines of evidence suggested that *Lnc956* KO caused severe DSBs and cytosolic dsDNA which induced innate immune as well as inflammation responses in post-implantation embryos. The inflammation can cause female-biased embryonic death after sex determination, whilst male embryos are less susceptible due to the anti-inflammatory effect of testosterone produced at high level in embryonic testes from approximately E12.5 onward ^32^. Based on the knowledge and our observation, we speculated that *Lnc956* KO embryos might die of lethal inflammation. We then administrated ibuprofen, an anti-inflammatory drug, in drinking water to pregnant females from gestation day 7.5 onward. Intriguingly, ibuprofen administration partially rescued the biased female embryo death (Fig. 3h), supporting the genomic instability and associated inflammation as the causes of embryonic lethility.

### *Lnc956*-TRIM28-HSP90B1 forms inter-dependent ribonucleoprotein (RNP) complex on replication forks under stress condition

*Lnc956* regulates CMG/replisome retention on chromatin and is critical for embryogenesis. To find out the underlying molecular mechanisms, we performed in vivo RNA pulldown combined with mass spectrometry analyses in order to identify its potential interaction proteins (Supplementary data 3). Meanwhile, we also performed iPOND followed by mass spectrometry to find proteins on replication forks of ESCs (Supplementary data 3). By comparing the two sets of protein list, we identified TRIM28 and HSP90B1 as overlapped candidates (Fig. 4a) and proposed that *Lnc956* might interact with TRIM28 and HSP90B1 and form a ribonucleoprotein (RNP) complex on replication forks. To test this hypothesis, we first examined the association of *Lnc956* with both proteins. Immunoblotting analysis of in vivo RNA pulldown samples showed that *Lnc956* interacted with TRIM28 and HSP90B1, and the association was stimulated by HU or aphidicolin treatment (Fig. 4b). Intriguingly, TRIM28 and HSP90B1 physically interacted and their association was robustly enhanced by HU (Fig. 4c) or aphidicolin (Fig. 4d). Of note, the stress-induced TRIM28-HSP90B1 interaction was dramatically suppressed by RNase A, implicating that RNAs mediated the protein interaction (Fig. 4c, d). We also validated the localization of TRIM28 and HSP90B1 on replication forks by iPOND assay (Fig. 4e). Importantly, the recruitment of two proteins to replication forks was dynamically regulated by replication stress. Initially, HU or aphidicolin treatment stimulated the accumulation of two proteins on replication forks. However, the levels of fork-resided proteins gradually decreased along with the replication stress treatment (Fig. 4f). This dynamics was different from the pattern observed for other components of replisomes including PCNA and RPA32 (Fig. 4f). The decrease could be due to the fact that prolonged replication stress continuously induced stalling of newly fired forks ^33^, and the increasing numbers of stalled forks gradually exhaust the available TRIM28-HSP90B1 pool, leading to the reduction of TRIM28-HSP90B1 level on each replication fork.

**Fig. 4.**
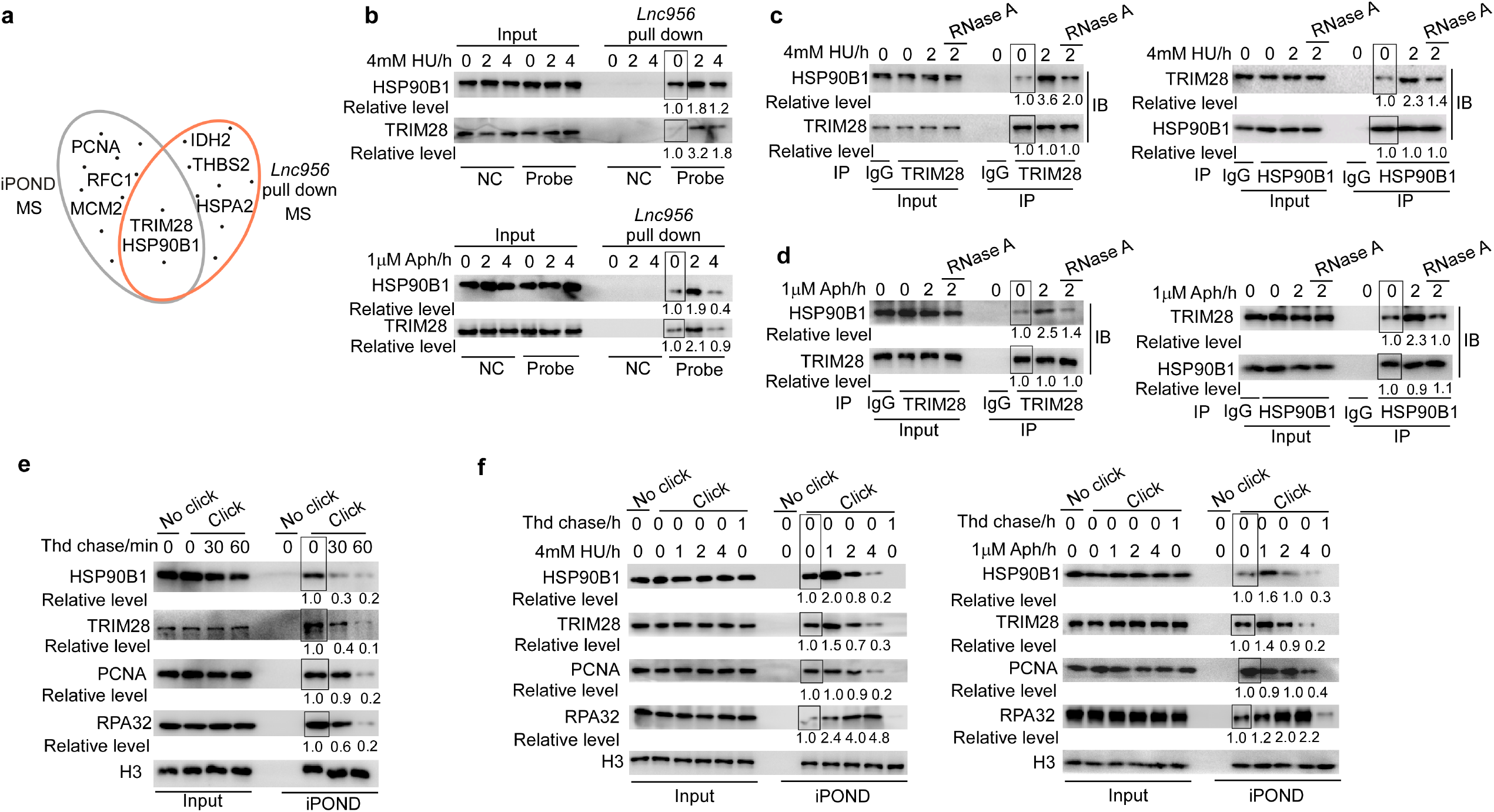
*Lnc956*-TRIM28-HSP90B1 forms ribonucleoprotein (RNP) complex on replication forks. **a** Mass spectrometry analyses identified TRIM28 and HSP90B1 as common candidates to reside on replication forks and interact with *Lnc956*. **b** In vivo RNA pull down validated the interaction of *Lnc956* with HSP90B1 and TRIM28. The interaction was enhanced by HU (upper panel) or aphidicolin (Aph) treatment (lower panel). Pulldown assay using sense probe was included as negative control (NC). **c-d** Reciprocal immunoprecipitation (IP) validated the physical interaction of HSP90B1 with TRIM28. The interaction was enhanced by HU (**c**) or aphidicolin (**d**) treatment and required the mediation of RNA species. **e** Immunoblotting of iPOND samples validated the localization of HSP90B1 and TRIM28 on replication forks of ESCs. Cells were chased with 10 μM thymidine chase (Thd) for different times. **f** The accumulation of HSP90B1 and TRIM28 on replication forks was dynamically regulated by HU (left panel) or aphidicolin treatment (right panel). The relative protein levels were normalized by input (b-d) or by histone H3 (e, f), and the levels in samples marked with box were set as 1. In (b-f), all experiments were repeated three times with similar results.

To gain more evidence supporting the formation of *Lnc956* -TRIM28-HSP90B1 RNP complex on replication forks, we examined the inter-dependence of the three components on replication forks. In *Lnc956* KO ESCs that we derived from mutant blastocysts (Supplementary Fig. 5a, b), the loading of TRIM28 and HSP90B1 on replication forks was not affected under unperturbed condition. However, their recruitment to stalling forks was impaired by the absence of *Lnc956* under HU treatment conditions (Fig. 5a). Concordantly, replication stress-induced TRIM28-HSP90B1 interaction was compromised by *Lnc956* KO (Fig. 5b). Re-expression of *Lnc956* in KO ESCs (Supplementary Fig. 5a, b) completely rescued these defects (Fig. 5a, b). Next we efficiently knocked down TRIM28 by two independent DOX-inducible shRNAs (Supplementary Fig. 5c) and examined its influences on HSP90B1 and *Lnc956*. TRIM28 KD had no obvious impact under normal condition. However, it decreased the loading of HSP90B1 (Fig. 5c, Supplementary Fig. 5d) and *Lnc956* (Fig. 5d, Supplementary Fig. 5e) on replication forks under HU treatment condition. Consistently, TRIM28 KD disrupted *Lnc956-*HSP90B1 association (Fig. 5e, Supplementary Fig. 5f). Likewise, HSP90B1 KD (Supplementary Fig. 5g) exhibited similar effects on TRIM28 and *Lnc956* under stressful condition (Fig. 5f-h, and Supplementary Fig. 5h-j). Taken together, these data suggest that *Lnc956*-TRIM28-HSP90B1 form a RNP complex on replication forks in an inter-dependent manner under replication stress condition. Assembly of this RNP complex facilitates accumulation of TRIM28 and HSP90B1 on replication forks. Notably, we found that this RNP assembly relied on the ATR replication checkpoint. Inhibition of ATR activation by specific inhibitor VE-821 compromised the accumulation of TRIM28, HSP90B1 (Fig. 5i) and *Lnc956* (Fig. 5j) on replication forks. TRIM28-HSP90B1 association was also compromised by VE-821 (Fig. 5k).

**Fig. 5.**
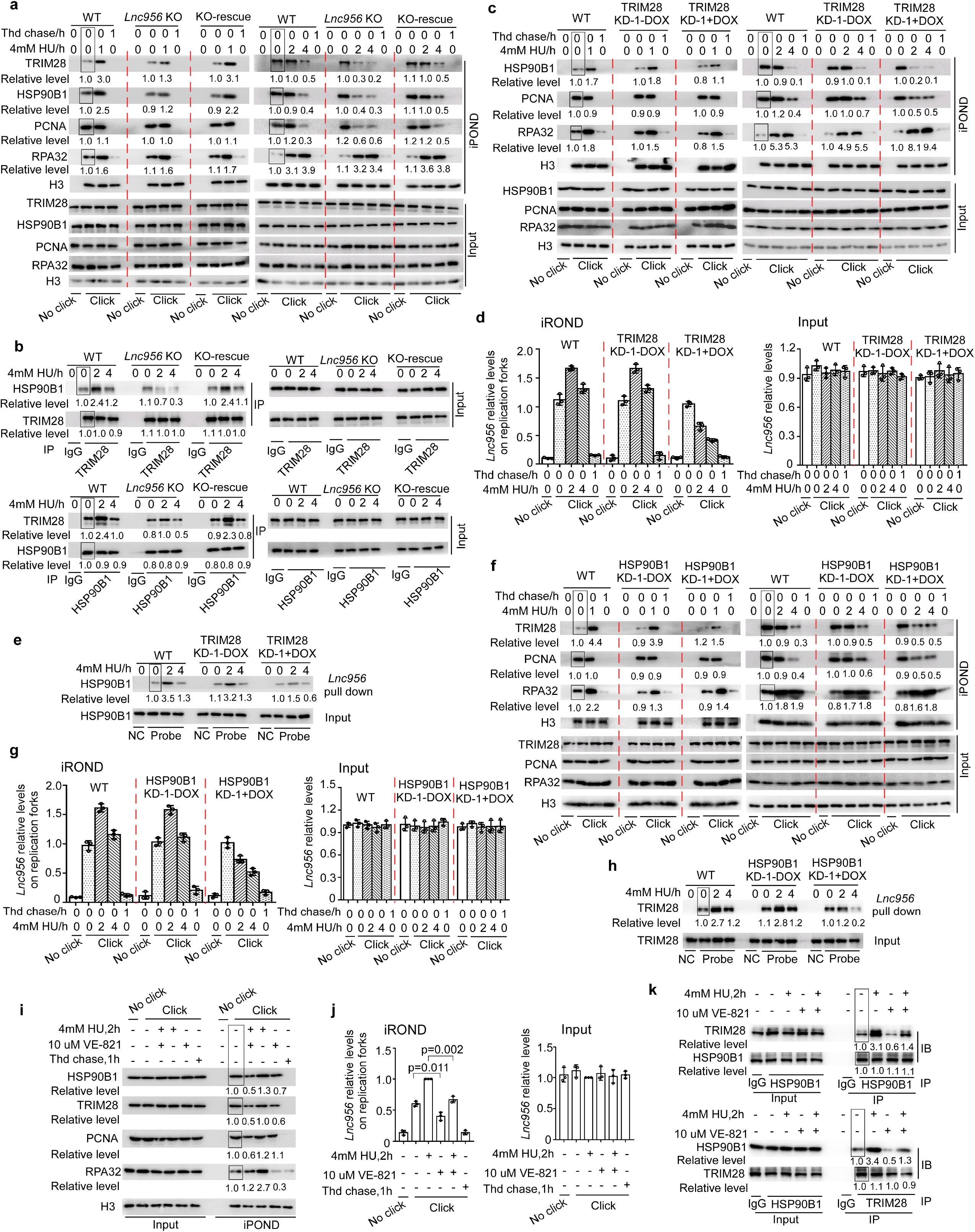
*Lnc956*-TRIM28-HSP90B1 RNP complex forms in an inter-dependent manner. **a** iPOND showed that *Lnc956* knock out (KO) impaired the loading of HSP90B1 and TRIM28 on stalled replication forks at all time-points of HU treatment. Re-expression of *Lnc956* (KO-rescue) could rescue the defects. **b** *Lnc956* KO also weakened the TRIM28-HSP90B1 interaction, which was restored by re-expression of *Lnc956*. **c** TRIM28 KD (KD-1) reduced the accumulation of HSP90B1 on stalled replication forks. **d** qRT-PCR analysis of iROND samples revealed the decreased accumulation of *Lnc956* on stalled forks in TRIM28 KD ESCs. The levels of *Lnc956* in inputs were normalized by *Actb*. **e** TRIM28 KD compromised the association of HSP90B1 with *Lnc956* as revealed by in vivo RNA pulldown. **f-h** Similarly, HSP90B1 KD (KD-1) decreased the accumulation of TRIM28 (**f**) and *Lnc956* (**g**) on stalled replication forks, and impaired the interaction of TRIM28 with *Lnc956* (**h**). **i-j** Inhibition of ATR activation by specific inhibitor VE-821 decreased the allocation of TRIM28 (**i**), HSP90B1 (**i**), and *Lnc956* (**j**) on replication forks. **k I**nhibition of ATR activity also decreased the TRIM28-HSP90B1 interaction. In vivo RNA pulldown using sense probe was set as negative control (NC) in (e, h). The protein levels were normalized by histone H3 in (a, c, f, i), and by input in (b, e, h, k). The relative protein levels in samples marked with box were set as 1. All experiments were repeated three times with consistent results.

Because *Lnc956*-TRIM28-HSP90B1 complex was formed in an inter-dependent manner, we speculated that loss of each component should exhibit similar phenotypes. Indeed, KD of TRIM28 or HSP90B1 accelerated the dissociation of replisome from chromatin, as indicated by the reduced levels of CMG components and PCNA on purified replication forks (Supplementary Fig. 6a, b) or on whole chromatin (Supplementary Fig. 6c, d). Consistently, stalled fork restart was compromised (Supplementary Fig. 7a, b) and inter-origin distances were increased (Supplementary Fig. 7c, d) in TRIM28 or HSP90B1 KD ESCs. Thus, *Lnc956*-TRIM28-HSP90B1 complex functions as a whole to regulate the CMG/replisome retention on chromatin under replication stress.

### *Lnc956*-TRIM28-HSP90B1 complex physically interacts with MCM2-7 hexamer

Next, we sought to understand how this RNP complex regulated CMG/replisome retention on chromatin. We first examined whether the RNP complex physically interacted with replisomes. Proteomic analysis of TRIM28 immunoprecipitates identified a long list of potential interaction proteins (Supplementary data 3). HSP90B1, the validated interaction partner, was in the list. Intriguingly, we detected all six components of MCM2-7 hexamer (MCM2, MCM3, MCM4, MCM5, MCM6 and MCM7), whereas the other components of active CMG complex, CDC45 and GINS, were not detected (Supplementary data 3). This suggested that *Lnc956*-TRIM28-HSP90B1 RNP complex might interact with MCM hexamer. We then went on to validate the association between the two complexes. MCM7 could pull down TRIM28 and HSP90B1 (Fig. 6a). Concordantly, TRIM28 or HSP90B1 co-precipitated with MCM7 but not CDC45 or GINS (Fig. 6b, c). In line with the induced formation of RNP complex under replication stress, RNP-MCM interaction was elevated after HU treatment (Fig. 6a-c). Thus, *Lnc956*-TRIM28-HSP90B1 RNP complex physically associated with MCM hexamer.

**Fig. 6.**
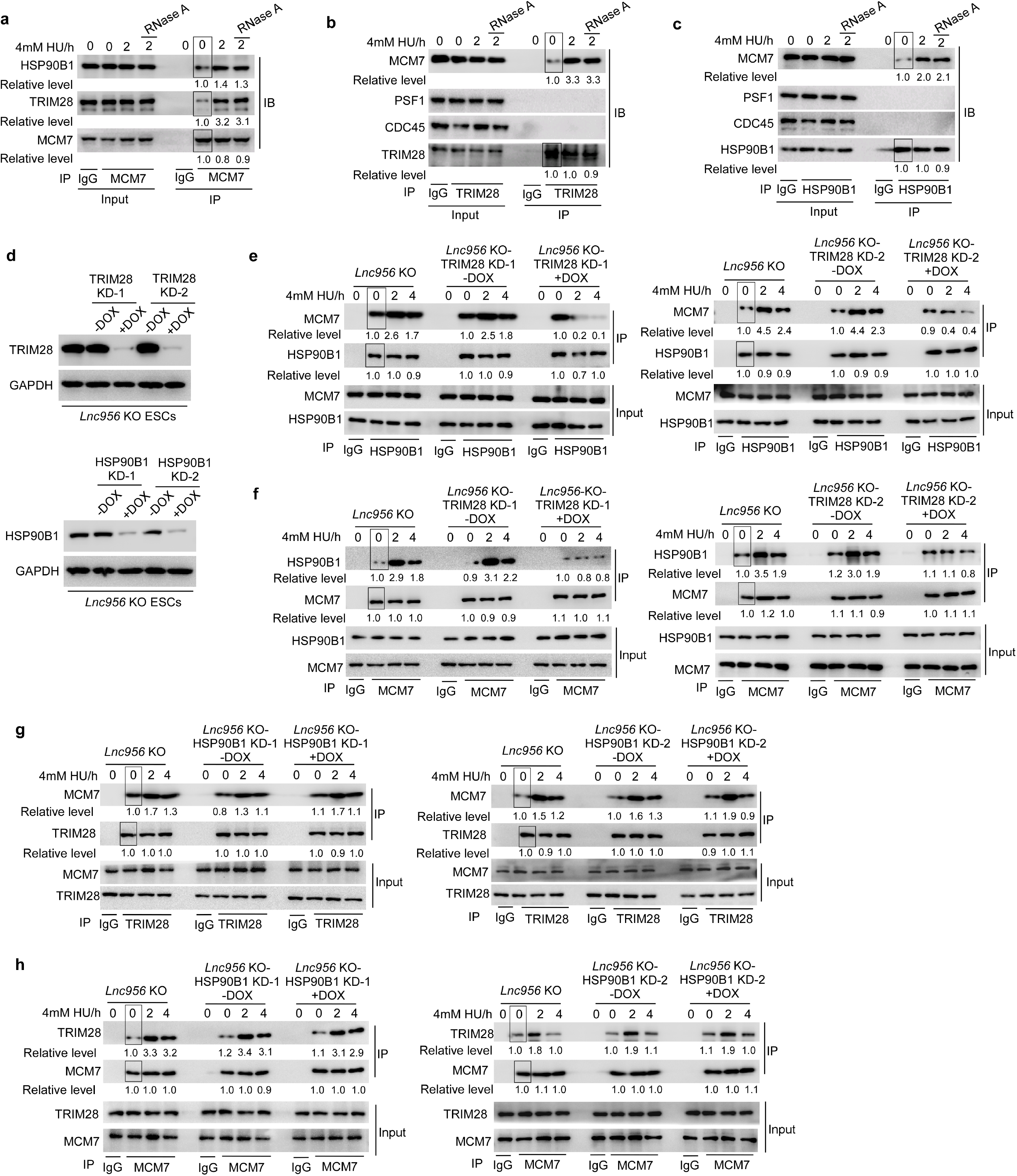
*Lnc956*-TRIM28-HSP90B1 RNP complex physically interacts with MCM hexamer. **a** MCM7 co-immunoprecipitated with TRIM28 and HSP90B1. The interaction was enhanced by HU treatment and did not rely on RNA species. **b-c** Reciprocally, TRIM28 (**b**) and HSP90B1 (**c**) could immunoprecipitate with MCM7, but not PSF1 or CDC45. The interaction was also stimulated by HU treatment and did not rely on RNAs. **d** KD of TRIM28 (upper panel) or HSP90B1 (lower panel) in *Lnc956* KO mESC using two independent DOX-inducible shRNAs. **e-f** Independent KD of TRIM28 using two shRNAs in *Lnc956* KO ESCs consistently impaired the association of HSP90B1 with MCM7, as revealed by reciprocal immunoprecipitation. **g-h** However, HSP90B1 KD using two shRNAs in *Lnc956* KO ESCs had no influence on the interaction between TRIM28 and MCM7, as shown by reciprocal immunoprecipitation. The relative protein levels were normalized by input, and the levels in samples marked with box were set as 1. All experiments were repeated three times with consistent results.

We then investigated which component of the RNP complex mediated the association. To dissect if *Lnc956* is necessary for RNP-MCM interaction, we depleted *Lnc956* by RNase A and examined its influence. The interaction between MCM7-TRIM28 or MCM7-HSP90B1 was not affected by RNase A treatment (Fig. 6a-c), indicating that *Lnc956* was dispensable for RNP-MCM association. This was consistent with the observation that none of MCM component was detected by mass spectrometry analyses in *Lnc956* in vivo pulldown assay (Supplementary data 3). To understand the role of TRIM28 and HSP90B1 in mediating RNP-MCM interaction, we individually depleted TRIM28 or HSP90B1 in *Lnc956* KO ESCs (Fig. 6d). Reciprocal immunoprecipitation showed that TRIM28 KD with two independent shRNAs severely impaired the association between MCM7 and HSP90B1 under replication stress condition (Fig. 6e, f). However, HSP90B1 KD did not affect MCM7-TRIM28 interaction (Fig. 6g, h). These data altogether supported that TRIM28 mediated the interaction between *Lnc956*-TRIM28-HSP90B1 RNP complex and MCM hexamer. A previous study reported the association of TRIM28 (also known as KAP1) with MCM and PCNA at heterochromatin regions ^34^. We wondered whether this RNP localized at heterochromatin. Counter-staining of HP1 (marker of heterochromatin) with *Lnc956* revealed partial co-localization (Supplementary Fig. 7e). Thus, the function of *Lnc956*-mediated RNP was not limited to heterochromatin regions.

### HSP90B1 regulates the retention of CMG helicase under replication stress

Unloading of CMG from chromatin is regulated by a series of sequential events. Initially, MCM7 of CMG complex undergoes K48-poly-ubiquitylation. Following ubiquitylation, the CMG complex is extracted from chromatin by segregase P97 ^35^. The ubiquitylation is prohibited by the DNA structure of active forks during elongation ^36^. Considering that the DNA structural change might regulate MCM7 ubiquitylation and CMG unloading, we speculated that chaperone protein HSP90B1, which functions as protein folding reservoir and stabilizes its clients ^37, 38^, might buffer the stress-induced changes and prevent the ubiquitylation reaction and CMG extraction by P97. To test this hypothesis, we used two different HSP90B1 inhibitors PU-WS13 and GRP94i ^39, 40^, to specifically block ATPase activity essential for its chaperoning function, and examined the influences on CMG retention, MCM7 ubiquitylation, fork breakage and stalling fork restart. We treated the WT ESCs with 4 mM HU for 2 h to induce modest replication stress, under which the CMG helicase was stably associated with chromatin. Inhibition of HSP90B1 activity by PU-WS13 at increasing concentrations did not affect the retention of TRIM28 and HSP90B1 on replication forks. However, it induced the premature unloading of CMG components from chromatin in a dose-dependent manner under mild stress condition. Intriguingly, PCNA, the other core component of replisome, remained stable on replication forks (Fig. 7a). Consistent results were obtained by using different inhibitor GRP-94i (Fig. 7b). Thus, inhibition of HSP90B1 activity specifically affected the CMG helicase and induced its unloading from chromatin. This defect was distinct to fork collapse-induce phenotype in which PCNA dissociates earilier than CMG helicase ^25^. We also investigated the influence of HSP90B1 activity inhibition on MCM7 ubiquitylation. Immunoblotting analysis revealed that MCM7 proteins, which were enriched by immunoprecipitation through MCM7 antibody, underwent K48- and K63-ubiquitylation after HU treatment and the modification level increased with the stress intensity ^41, 42^. Strikingly, inhibition of HSP90B1 activity with either inhibitor sharply increased both types of ubiquitylation levels (Fig. 7c), indicating that HSP90B1 attenuated the MCM7 ubiquitylation. To further clarify the involvement of ubiquitylation-P97 pathway in the extraction of CMG, we blocked P97 activity in the presence of HSP90B1 inhibitor, and examined if it could prevent the extraction of CMG. P97 inhibitor NMS-873 or CB-5083 ^43, 44^ did not affect the MCM7 ubiquitylation in ESCs treated with HU plus HSP90B1 inhibitor PU-WS13 (Fig. 7d). However, both P97 inhibitors prevented the unloading of CMG in a dose-dependent manner, as revealed by iPOND (Fig. 7e) and chromatin purification analyses (Fig. 7f). These results altogether suggested that HSP90B1 promoted the retention of CMG helicase probably through blocking the MCM7 ubiquitylation-P97 extraction pathway.

**Fig. 7.**
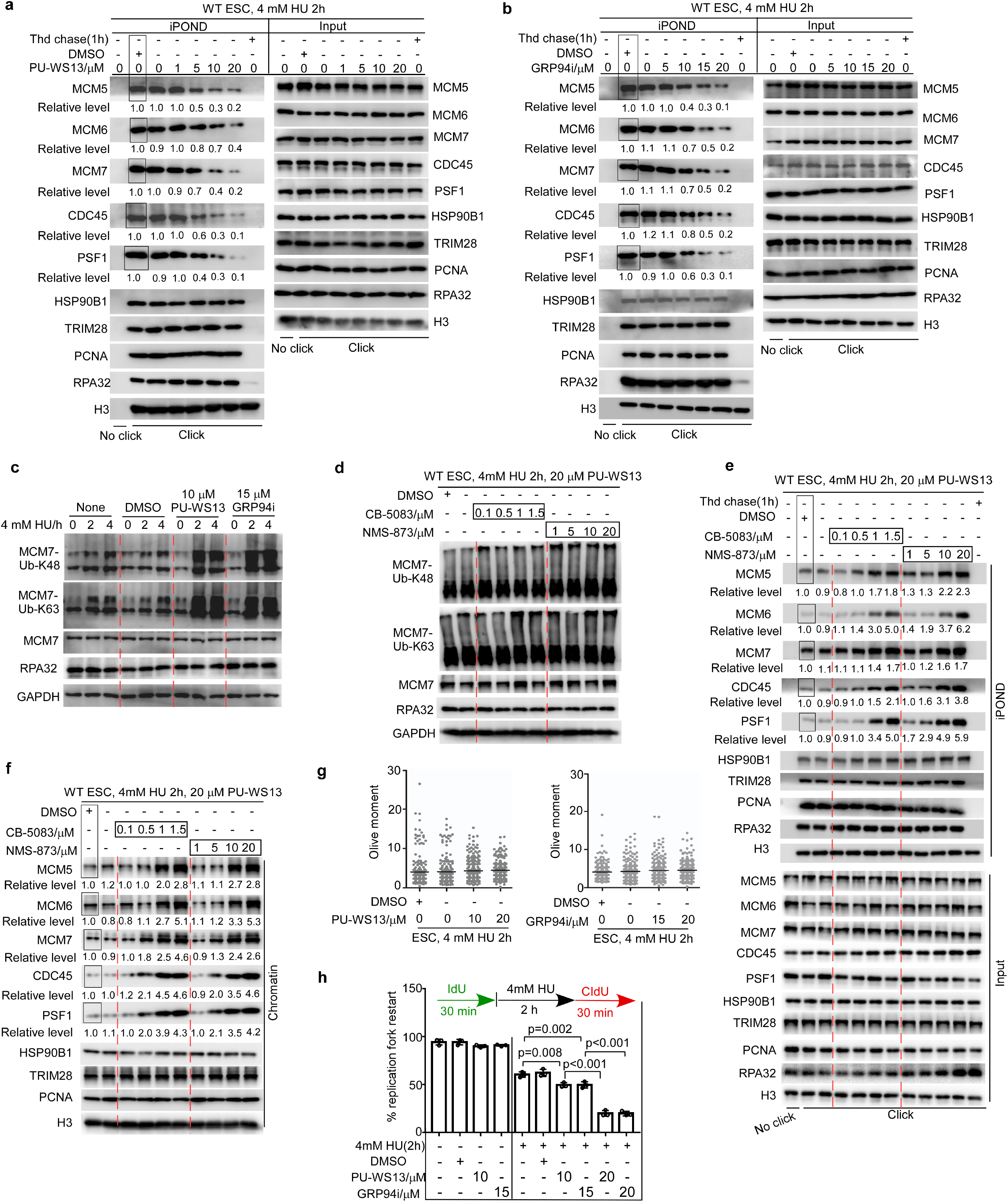
HSP90B1 stabilizes CMG helicase under replication stress. **a-b** ESCs were treated with 4 mM HU for 2 h to induce mild replication stress. Under this condition, inhibition of HSP90B1 ATPase activity by inhibitor PU-WS13 (**a**) or GRP94i (**b**) consistently induced the dissociation of CMG helicase from stalled replication forks in a dose-dependent manner. In contrast, fork-associated HSP90B1, TRIM28, PCNA and RPA32 were not affected. **c** Inhibition of HSP90B1 activity by PU-WS13 and GRP94i increased the K48 and K63 ubiquitynation of MCM7 under the HU treatment condition. **d** ESCs were treated with HU plus PU-WS13 to induce K48 and K63 ubiquitynation of MCM7. Under this condition, blocking P97 activity by inhibitor CB-5083 or NMS-873 did not affect the MCM7 ubiquitylation. **e** However, blockage of P97 activity prevented CMG unloading and increased CMG retention on replication forks. **f** Consistent results were obtained when whole chromatin was analyzed. The relative protein levels were normalized by histone H3 in (a, b, e, f). The levels in samples marked with box were set as 1. All experiments were repeated three times with similar results. **g** Neutral comet assay showed that inhibition of HSP90B1 activity by PU-WS13 (left panel) or GRP94i (right panel) under HU treatment condition had no influence on the formation of DNA double strand breaks (DSBs). At least 200 tails from three replications were analyzed in each group. **h** Inhibition of HSP90B1 activity by PU-WS13 or GRP94i impaired the stalled fork restart in a dose-dependent manner. At least 200 fibers from three independent experiments were analyzed. Data were shown as mean ± SEM, two-tailed Student’s *t*-test.

Next, we examined the effect of HSP90B1 activity inhibition on the dynamics of fork breakage or DSB formation. Intriguingly, under the same treatment conditions, HSP90B1 inhibition didn’t induce fork breakage and DSB formation (Fig. 7g). This observation was similar to that in *Lnc956* KD ESCs, further supporting that fork breakage and replisome unloading were separately regulated in replication stress response. Because CMG helicase can not be reloaded once unloaded during S phase, we speculated that stalled fork restart was compromised by HSP90B1 inhibition even the fork remained intact. Indeed, inhibition of HSP90B1 activity by either inhibitor impaired the replication fork restart in a dose-dependent manner following replication stress (Fig. 7h). Taken together, these data supported that fork breakage and replisome unloading were independently regulated. *Lnc956*-TRIM28-HSP90B1 RNP complex directly regulated CMG retention and this function was largely mediated by HSP90B1. In addition, dissociation of CMG helicase from chromatin did not influence TRIM28-HSP90B1 on replication forks (Fig. 7a, b), suggesting that assembly of the RNP complex (*Lnc956*-TRIM28-HSP90B1) on replication forks did not rely on CMG helicase.

### Ectopic expression of *Lnc956* enhances CMG/replisome stability in somatic cells

Depletion of *Lnc956* in mouse ESCs revealed its essential role in driving the assembly of a novel RNP complex, which associates with MCM hexamer and regulates CMG retention on chromatin under stress condition. To further illustrate the functional significance of *Lnc956*, we ectopically expressed *Lnc956* in NIH3T3 cells (Supplementary Fig. 7f) and investigated the effects. As shown, expression of *Lnc956* in NIH3T3 increased the fork-allocated TRIM28 and HSP90B1 levels (Fig. 8a). Concordantly, TRIM28-HSP90B1 interaction was enhanced by *Lnc956* overexpression (Fig. 8b). As a result, *Lnc956* expression promoted the CMG/replisome retention on chromatin (Fig. 8a, c) and the fork restart ability (Fig. 8d) in response to HU treatment. Notably, the dormant origin density was also increased (Fig. 8e), reflecting the increased retention of MCM hexamers on chromatin. Blocking the ATPase activity of HSP90B1 counteracted the beneficial effects of *Lnc956* on MCM retention, stalled fork restart, as well as the dormant origin density in NIH3T3 cells (Fig. 8d-f). These results supported that introduction of *Lnc956* in somatic cells was able to drive the assembly of regulatory RNP complex on replication forks, albeit with low efficiency. Other unknown ESC-specific mechanisms could be required for its efficient assembly. In summary, we reported a novel RNP complex composed of *Lnc956*-TRIM28-HSP90B1 and assembled under replication stress condition. This RNP complex is physically associated with MCM hexamer via TRIM28 and regulated MCM2-7 retention on chromatin via HSP90B1 (Fig. 8g). The formation of this RNP complex significantly increases the local concentrations of TRIM28 and HSP90B1, confering cells with greater ability to retain CMG helicase on chromatin and to ensure genomic stability.

**Fig. 8.**
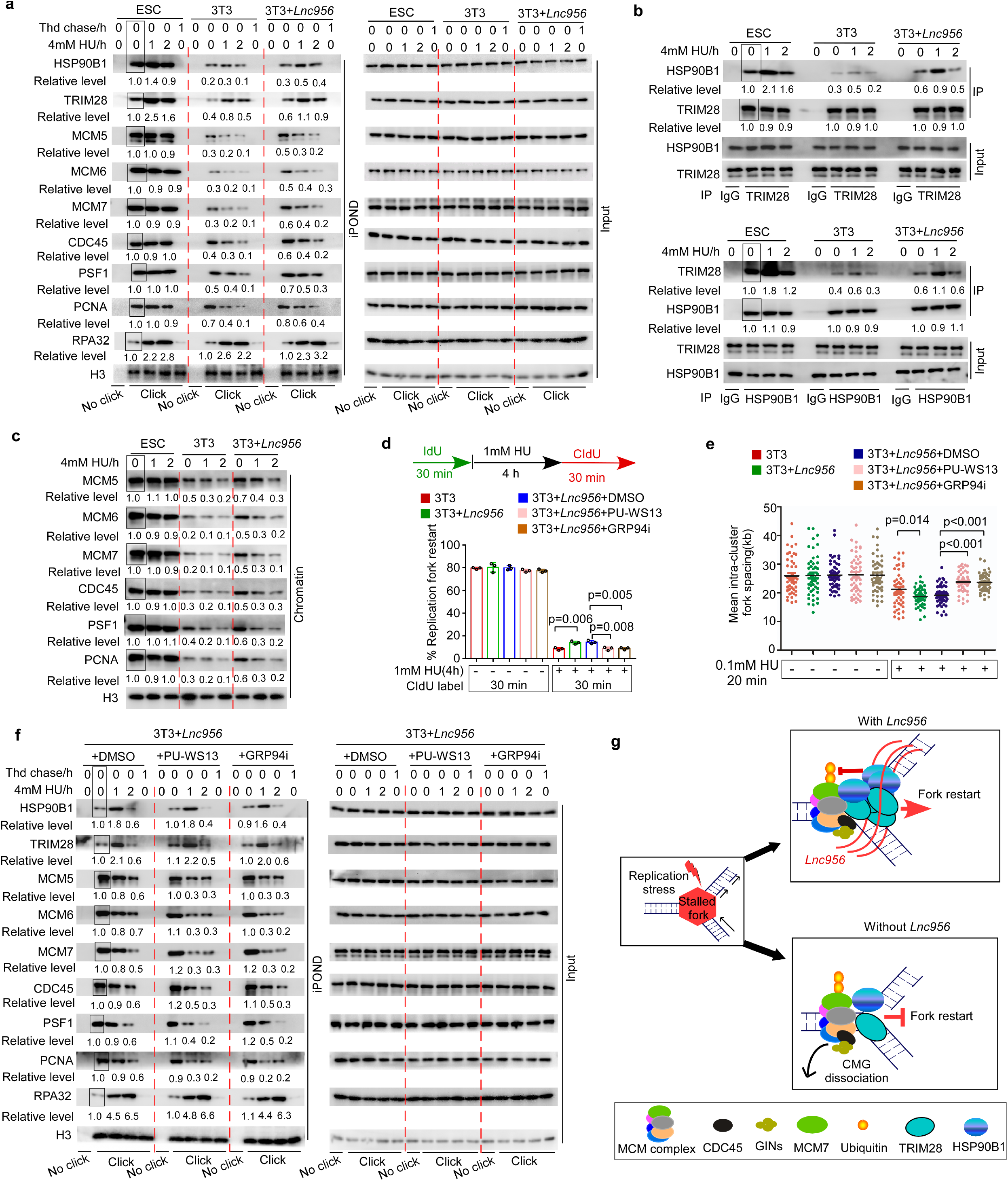
Ectopic expression of *Lnc956* in somatic cells enhances CMG retention on chromatin. **a** iPOND analysis revealed that ectopic expression of *Lnc956* in NIH3T3 cells increased the fork-resided TRIM28 and HSP90B1 levels, and promoted the retention of CMG helicase on replication forks when compared to 3T3 cells. **b** Reciprocal immunoprecipitation showed that ectopic expression of *Lnc956* in NIH3T3 cells enhanced TRIM28-HSP90B1 interaction. **c** Immunoblotting analysis of purified chromatin showed that ectopic expression of *Lnc956* in NIH3T3 cells increased CMG association with chromatin. **d** Ectopic expression of *Lnc956* in NIH3T3 cells promoted the stalled fork restart. This beneficial effect was counteracted by inhibition of HSP90B1 activity. At least 200 DNA fibers from three independent experiments were analyzed. **e** Ectopic expression of *Lnc956* in NIH3T3 cells increased dormant origin density under replication stress, and this effect was counteracted by HSP90B1 inhibitors. At least 50 fibers from three independent experiments were analyzed. **f** Inhibition of HSP90B1 activity by PU-WS13 or GRP94i did not influence the accumulation of HSP90B1 and TRIM28 on replication forks. However, it counteracted the beneficial effect of *Lnc956* on CMG retention on chromatin. **g** The working model of *Lnc956*-TRIM28-HSP90B1 RNP complex on replication forks. The relative protein levels were normalized by histone H3 in (a, c, f) or by input in (b). The levels in samples marked with box were set as 1. All experiments were repeated three times with similar results. Data were shown as mean ± SEM, two-tailed Student’s *t*-test.

## Discussion

Replication stress is the major source of endogenous DNA damages. Dysregulation of replication stress response can cause variable human diseases ranging from developmental defects to cancers ^15, 45^. Numerous efforts have been made to understand the local and global cellular responses to stalled forks ^17^. Owing to the innovative methods such as iPOND, an increasing numbers of proteins were identified on replication forks under normal and/or stressed conditions. These regulatory proteins play critical roles in ensuring fork intact and restart under stressed conditions ^17, 46^. To our knowledge, it remains unexplored whether there exist non-coding RNAs as regulators on replication forks. LncRNAs are able to enhance the functions of their interacting protein by driving efficient protein condensation ^26, 27^. Using PSC as a model, we reported the presence of functional lncRNA on stalled replication forks and revealed its critical role in directly regulating CMG helicase retention to promote stalled fork restart. Importantly, our findings demonstrated that replication checkpoint directly regulates replisome retention on chromatin, which underlies fork collapse in PSCs. Stable retention of CMG helicase, which once unloaded from chromatin at S phase can not be reloaded, confers PSCs superior abilities to restart stalled forks and preserve genomic stability. Finally, we identified a novel lncRNA *Lnc956* essential for embryogenesis and provided new clue to decipher the cause of recurrent pregnancy loss in clinic.

*Lnc956* drives the *Lnc956*-TRIM28-HSP90B1 RNP complex assembly in an inter-dependent manner upon replication stress. A line of evidence from iPOND, iROND and RNA-FISH confirmed that this RNP complex was localized on stalled replication forks and associated with MCM hexamer. *Lnc956* depletion had no influence on the activation of ATR signaling, which is essential to evoke the global response to replication stress. Intriguingly, *Lnc956* depletion in ESCs or ectopic expression in somatic cells affected the density of dormant origins. In addition, *Lnc956* depletion caused robust (more than half) loss of MCM components from chromatin after the initial two hours of HU treatment. Because most chromatin-associated MCM hexamers exist in pre-replication complexes (pre-RCs), the drastic loss of MCM from chromatin could not be explained by the unloading of MCM complexes from active replication forks. Based on these observations, we hypothesized that the RNP complex might also bind to MCM hexamer of dormant origins and exhibit regulation on genome-wide MCM hexamers.

Assembly of *Lnc956*-TRIM28-HSP90B1 RNP complex significantly enhanced the recruitment of TRIM28 and HSP90B1 to stalled forks. Previous studies demonstrated that lncRNAs (for instance *NORAD*, *Xist*, *Neat1*) could drive the condensation and phase separation of interacting RNA binding proteins (RBP), thereby drastically increasing the local concentration and probably the activity of the RBPs ^27, 47, 48^. In the future, it would be intriguing to clarify whether *Lnc956* stimulates the accumulation of TRIM28 and HSP90B1 through phase separation. *Lnc956*-TRIM28-HSP90B1 RNP complex promotes CMG retention under replication stress. A line of evidence supported that this regulation was direct, rather than an indirect consequence of fork breakage prevention. First, CMG helicase unloading induced by *Lnc956* depletion or HSP90B1 inhibition occurred independent of the induction of fork breakage and DSB formation. Second, HSP90B1 inhibition did not affect PCNA association with nascent DNA, but specifically induced CMG components unloading. This did not comply with the nucleotide depletion-induced fork collapse in which PCNA is rapidly dissociated from nascent DNA, whereas little dissociation of CMG complex is detected until late time point of treatment ^17, 25^. Third, *Lnc956*-TRIM28-HSP90B1 RNP physically associates with MCM hexamer, providing a basis for direct regulation of RNP on CMG helicase. Finally, *Lnc956* depletion in mESCs or ectopic expression in somatic cells impacted the inter-origin distance, reflecting its influence on MCM hexamer retention in dormant origins. Previous studies suggested that the major functions of ATR replication checkpoint were to repair stalled forks, prevent fork cleavage, and modulate replisome function, rather than regulating replisome stability ^25, 49, 50^. To date, it still remains unclear whether replisome unloading is simply the outcome of fork collapse, or underlies fork collapse ^17^. The *Lnc956*-TRIM28-HSP90B1 RNP was assembled under the control of ATR signaling, therefore being a novel component of replication checkpoint. Our work suggested that the ATR replication checkpoint could also actively regulate replisome stability, which underlies fork collapse in certain celluar context.

We noticed that disruption of *Lnc956*-TRIM28-HSP90B1 RNP assembly by knocking down either component caused CMG and PCNA dissocation, whereas inhibition of HSP90B1 activity selectively destabilize CMG without affecting PCNA. This observation implicated that this RNP regulated replisome retention by distinct pathways. One pathway was mediated by HSP90B1, which probably utilized its ATPase activity to buffer the stress-induced changes and suppress MCM7 ubiquitylation required for MCM hexamer extraction by p97. The other pathway might be mediated by TRIM28. Previous studies showed that TRIM28 interacted with PCNA and functioned as SUMO E3 ligase to modify PCNA in resolving transcription-replication conflict ^51^. Our proteomic analysis of TRIM28 interacting partners also identified PCNA as one of interacting proteins. Whether TRIM28 interacts with PCNA and how TRIM28 regulates PCNA stability in ESCs require further investigation.

In summary, we identified a new lncRNA-based mechanism which is a novel component of ATR replication checkpoint and directly regulates the retention of CMG to prevent stalled fork collapse. This mechanism plays essential roles in embryogenesis. Future works are warranted to investigate the human *Lnc956* expressions and functions, and its relevance to human developmental abnormalities.

## Methods

### Generation of *Lnc956* knockout mice

The protocols of gene knockout, animal care and manipulation were approved by the Institutional Animal Care and Use Committee of the Kunming Institute of Zoology, Chinese Academy of Sciences. The *Lnc956^+/−^* C57Bl/6J mice were generated by Cyagen Company (https://www.cyagen.com) using CRISPR-based gene editing technology. The guide RNAs (gRNA) were designed based on mouse *Lnc956* genome region sequence (chr17 qA3.3: 28742231-28744321). Cas9 mRNA and a pair of gRNAs (5’-AGCGGGCAGCGCCCGGGAGGAGG-3’; 5’-TTGGAAGAGTAACACGTTTTGGG-3’) were co-injected into one-cell embryos to generate targeted knockout mice. The genome DNA from mouse tails was used for genotype confirmation by PCR amplification and DNA sequence analysis. The PCR program was: 94°C for 5 minutes, (94°C for 15 s, 60°C for 30 s, 72°C for 30 s) 35 cycles, 72°C for 15 minutes. Genotyping primers were shown in supplementary Table 1.

### Cell culture

The mouse embryonic fibroblasts (MEFs) were obtained from embryos of CD1 mice at embryonic day 13.5 (E13.5). Mitomycin-treated MEFs were used as co-culture feeder cells for derivation and culture of mouse embryonic stem cells (mESCs). Wide type and *Lnc956^-/-^* mESCs were derived from the inner cell mass of E3.5 blastocysts.

The culture of MEFs, NIH3T3 and mESCs were described in detail in our previous papers ^14^. Briefly, MEFs and NIH3T3 cells were maintained in DMEM minimal medium (Gibco, 11965) with 10% FBS (Gibco, 10099141C). mESCs were cultured in DMEM/F12 medium supplemented with 20% Knockout serum replacement (Gibco, 10828028), 2 mM L-glutamine (Sigma, G8540), 1mM sodium pyruvate (Gibco, 11360070), 0.1 mM β-mercaptoethanol (Sigma; M7522), 1% non-essential amino acids (Gibco, 11140-035) and 1000 units/mL mouse leukemia inhibitory factor (LIF) (Millipore, ESG1107). All cells were grown at 37°C and 5% CO_2_ atmosphere. Regular mycoplasma testing was performed using the LookOut Mycoplasma PCR detection (Sigma, Cat. No. MP0035).

### Lentivirus-mediated gene manipulation

The shRNA sequences and primers were listed in supplementary Table 2. Following the manufacturer’s instructions (Open Biosystems, QCHENG BIO, QCP0703), all the shRNA sequences were constructed into pTRIPZ lentivirus tet-on inducible shRNAmir system. The *Lnc956* and *Trim28* coding regions (CDS) were inserted into pTOMO-IRES-EGFP lentiviral expression vector (Addgene, #26291). *Lnc956* ORF1, ORF2, ORF3 and *Floped* CDS containing Flag-tag were inserted into pcDNA3.1 vector. For the virus package, the shRNAmir or pTOMO-IRES-EGFP lentiviral expression vectors were mixed with packaging plasmids psPAX2 and PMD2.G (shRNAmir or pTOMO-IRES-EGFP: psPAX2: PMD2.G=2:1:1) and co-transfected into 293T cells. For the groups of shRNAmir-mediated gene knockdown, 48 hours after lentivirus infection, the puromycin (0.5 μg/mL) was added into culture medium for selection. For the groups of pTOMO-IRES-EGFP mediated gene expression, GFP positive cells were sorted by flow cytometry for extended culture and analysis.

### Cell cycle

For cell cycle analysis experiment, the cells were digested with trypsin, washed with PBS once, and then immobilized overnight with pre-cooled 70% alcohol at 4℃. The fixed cells were washed once with PBS, then suspended in PBS containing 50 µg/mL PI (sigma, P4864-10ML) and 20 µg/mL RNase A, and incubated at 37℃ for 30 min. The DNA content was assessed by FACS LSRFortessa flow cytometry (BD). The raw data were analyzed using FlowJo™ software (version 7.6) and the Dean-jett-Fox algorithm.

### Isolate proteins on nascent DNA (iPOND)

iPOND was performed as described in previous paper ^19^. In order to harvest enough S phase cells, NIH3T3 cells were synchronized by 2 mM thymidine (sigma, T1895) for 15 hours, followed by fresh medium culture for 2.5 hours. For each sample, 10^8^ synchronized NIH3T3 or normal cultured mESCs were incubated with 10 mM EdU (Life Technologies, A10044) for 10 minutes, followed by PBS wash and hydroxyurea (HU) treatment. 10 μM thymidine chase was included as chromatin control. Next, the cells were fixed with 1% formaldehyde (Sigma, F1635) and then quenched with 0.125 M glycine (Sangon Biotech, A100167). The cells were scraped, and permeabilized by ice-cold 0.25% Triton X-100/PBS for 30 minutes at room temperature. After washing with 0.5% BSA/PBS, the cells were re-suspended in click reaction buffer including 10 mM Sodium ascorbate, 2 mM CuSO_4_, 10 μM Biotin-azide (Life technology, B10184) or DMSO (negative control) for 1 hour at room temperature. After the click reaction, cells were washed with 0.5% BSA/PBS for three times, and re-suspended in lysis buffer (50 mM Tris-HCl, pH 8.0; 1% SDS; 1 μg/mL aprotinin; 1 μg/mL leupeptin) for sonication (30 seconds pulse/30 seconds pause, 60 cycles) using BioruptorTM UCD-200 machine. Finally, samples were centrifuged at 10000×g for 10 minutes at 4°C to remove the precipitate. 1/1000 of each sample (volume ratio) was collected as an input sample. The remaining samples were incubated with streptavidin magnetic beads (Millipore, 69203) to purify replicative DNA fragments. 2×SDS Laemmli sample buffer (0.4 g SDS, 2 mL 100% Glycerol, 1.25 mL 1 M Tris pH 6.8, and 0.01 g Bromophenol blue in 8 mL H_2_O) was used to elute nascent DNA binding proteins. The eluted samples were boiled at 100°C for 10 minutes and used for western blotting analysis or mass spectrometry screening.

### Isolate RNAs on nascent DNA (iROND)

In order to purify potential RNA components on replication forks, we modified the iPOND protocol described above and developed iROND protocol. The basic principles and general procedures of iROND are the same as iPOND. To avoid RNA degradation, all the buffers and reagents were prepared using DEPC-treated ddH2O and 10 U/μL RNase inhibitors (Invitrogen, EO0382) in the whole procedure, and all the procedures were performed on ice or 4°C whenever possible. 500 μL Trizol reagent (Tiangen, DP424) was used for RNA purification of each sample, and RNA samples were used for RNA sequencing or qRT-PCR analysis. The quality control, libraries building and sequencing were performed by biolinker technology (kunming) co., ltd.

### Immunoblotting

Cells were lysed with RIPA buffer (Beyotime, P0013J). The protein lysate was electrophoresed into 6-10% SDS-PAGE gels and transferred to a PVDF membrane (Roche, 03010040001). After blocking with 5% BSA, the membranes were incubated with primary antibodies overnight at 4°C. After washing with TBS-T buffer for three times, the membranes were labeled with the corresponding HRP-conjugated secondary antibodies at room temperature for 1 hour. After washing with TBS-T buffer for three times, images were collected from the ProteinSimple FluorChem system (protein simple, Fluorchem M FM0561) after incubation with SuperSignal West Pico PLUS Chemiluminescent Substrate (Thermo Fisher Scientific, Cat. No. 34580). All antibody information is listed in Supplementary Table 3.

### RNA sequencing and analysis

The iROND samples and embryonic RNA samples were reverse transcribed into cDNA libraries using the TruSeqTM RNA Sample Preparation Kit (Illumina). RNA sequencing was performed with an Illumina HiSeq™ 3000 or HiSeq X Ten platform. The clean reads of mESC iROND samples were processed with Tophat2 and Cufflinks. Non-coding transcripts were annotated with NONCODE database (http://www.noncode.org/) (version 4.0). The Coding-Non-Coding Index (CNCI) software was used to identify novel lncRNAs. DEGseq software was used to analyze differentially expressed lncRNAs.

The clean reads of embryonic RNA samples were mapped to the mouse reference genome (MM10) using Tophat2 software and coded gene expression was calculated using Cufflinks. Cuffdiff software to determine differentially expressed genes, online tool (http://geneontology.org/) to enrich geneontology, with default parameters of ‘gplot ’r package to create heat maps. RNA-seq data have been deposited in Gene Expression Omnibus (GEO) database (login number :GSE196639).

### Real-time quantitative reverse transcription PCR (qRT-PCR)

Trizol (Tiangen, DP424) was used to extract total RNA from the cultured cells. The quality and concentration of the RNA samples was determined by spectrophotometry (Nanodrop, ND1000). 1 g RNA was subjected to DNase treatment and reverse transcribed into cDNA. TB Green™ Premix Ex Taq™ II kit (Takara, RR820A) and CFX96TM-Real Time Systerm (Biorad, CFX96Touch) were used for real-time quantitative analysis. All reactions were normalized by *Actb*. The primers were provided in Supplementary Table 1.

### RNA fluorescence in situ hybridization (RNA-FISH) and Immunofluorescence staining

RNA-FISH was performed as described in published paper ^52^. All operations were performed in RNase free conditions. The cells were cultured on a glass lid (ZEHAO, 12-545-82) pre-coated with 3 mg/mL Matrigel. In order to label replication forks, 10 µM BrdU was added into culture medium for 10 minutes. Cells were then fixed with 4% paraformaldehyde for 10 minutes, followed by 0.2% Triton X-100/PBS treatment for 5 minutes at room temperature. Cells were hybridized with the 2 ng DNA probe (Guangzhou Ribo Biological Co., Ltd.) at 37°C overnight, then washed for 3 times with 4×Sodium Citrate Buffer (SSC), 1 time with 2×SSC, 1 time with 1×SSC at 42°C and 1 time with PBS at room temperature.

For the co-labeling of proteins and BrdU, the slides were re-fixed with 4% paraformaldehyde for 10 minutes at room temperature, denatured with 2.5 M HCL for 30 minutes at room temperature, followed by neutralization with 0.1 M sodium borate for 10 minutes. Then, the slides were blocked with 2% BSA for 1 hour, and incubated with the primary antibodies overnight at 4°C. After washing 3 times with 0.2% Tween 20/PBS, cells were incubated with fluorescence-conjugated secondary antibodies for 1 hour at room temperature. DAPI was counterstained for 10 minutes. Slides were sealed with glycerin. Images were captured using confocol microscope (Olympus, FV1000).

### Rapid amplification of cDNA ends (RACE)

The RACE was performed according to published protocol ^53^. Briefly, the template switching oligonucleotides (TSO), oligo-dT primers and ISPCR primers were ordered as Smart-seq2 protocol. To obtain the 5’ end of *Lnc956*, gene-specific primer 1 (GSP1) was used to reverse transcription, template switching was performed with TSO primers, gene-specific primer 1 and ISPCR primer were used to amplify the 5’ end by PCR. To obtain the 3’ end of *Lnc956*, oligo-dT primers was used to reverse transcription, gene-specific primer 2 (GSP2) and ISPCR primers were used to amplify the 3’ end by PCR. The products were constructed into the pEasy-blunt vector and verified by sequencing. The primers were provided in Supplementary Table 1.

### Subcellular fractionation

2 × 10^7^ cells were lysed in 200 µL cold cytoplasmic lysis buffer (0.15% NP-40, 10 mM Tris pH 7.5, 150 mM NaCl) for 5 minutes, added 500 µL sucrose buffer (10 mM Tris pH 7.5, 150 mM NaCl, 24% sucrose) and centrifuged at 13000×g for 10 minutes, The supernatant was collected as cytoplasmic fraction. The pellet was washed with 200 µL cytoplasmic lysis buffer without NP-40 and resuspended in 200 µL ice cold glycerol buffer (20 mM Tris pH7.5, 75 mM NaCl, 0.5 mM EDTA, 50% glycerol, 0.85 mM DTT) and 200 µL ice cold nuclei lysis buffer (20 mM HEPES pH 7.5, 7.5 mM MgCl_2_, 0.2 mM EDTA, 0.3 M NaCl, 1 M Urea, 1% NP-40, 1 mM DTT), vortexed and incubated on ice for 1 minute. After centrifuge at 14000×g for 2 minutes, the supernatant was collected as nucleoplasm fraction. The pellet was resuspended in 50 µL PBS as chromatin fraction. The separated fractions were mixed with 2×SDS Laemmli sample buffer for immunoblotting, or mixed with Trizol reagent for RNA extraction and qRT-PCR analysis. To avoid RNA degradation, all the buffers and reagents were added with 10 U/μL RNase inhibitors (Invitrogen, EO0382) in the whole procedures.

### Protein ubiquitynation assay

10^6^ cells were harvested in 150 μL SDS lysis buffer (50 mM Tris–HCl, pH 6.8, 1.5% SDS). The samples were boiled at 100 °C for 15 min. 100μL of protein lysate was diluted with 1.2 mL EBC/bovine serum albumin (BSA) buffer (50 mM Tris–HCl, pH 6.8, 180 mM NaCl, 0.5% CA630, 0.5% BSA) and incubated with protein G beads (Sigma, 16-662) and antibody overnight at 4 °C with rotation. The beads were collected by centrifugation at 5 000 g for 3 min at 4 °C and washed 5 times with 1 mL ice-cold EBC/BSA buffer. Proteins were resuspended with 30 μL of SDS sample loading buffer and analyzed by immunoblotting.

### Dot blotting detection of cytosolic double strand DNA (dsDNA)

E12.5 embryos were dissociated into single cells with collagenase (Sigma, C6745) at 37 °C. Cells were used for downstream subcellular fractionation. 2 µL cytoplasmic fraction was dropped onto nylon membrane (Sigma, 15356) to immunoblotting, and nuclear fractionation was used as control.

### DNA fiber assay

DNA fiber analysis was performed as described ^14, 29^. 50 µM 5-iodo-2’-deoxyuridine (IdU; Sigma, I7125) was used to label the cells for 30 minutes, with or without hydroxyurea (HU, Selleck, S1896) treatment, then 250 µM 5-chloro-2’-deoxyuridine (CldU; Sigma, C6891) was used for the second labeling. After labeling, ∼3000 cells in 2.5 µL suspension were dropped onto one end of the glass slide, mixed with 7.5 µL lysis buffer (50mM EDTA, 0.5%SDS, 200mMTris-HCl, pH7.5), incubated for 8 minutes at room temperature. Then, the slides were tilted to 15° to spread the DNA fibers along the slide. The slides were then treated with 2.5 M hydrochloric acid, incubated with rat anti-BrdU/CIdU (BU1/75) monoclonal antibody (Novus, NB500-169) and mouse anti-IdU monoclonal antibody (BD, 347580). The secondary antibodies were AlexaFluor Cy3-conjugated goat anti-rat and AlexaFluor 488-conjugated goat anti-mouse respectively. Images were captured using confocol microscope (Olympus, FV1000). Image J software was used to measure the length of DNA fibers, and the formula that 1 μm=2.59 kb was used to convert the μm value to kb. New replication origin firing analysis was performed as described ^54^. Cells were labeled with CIdU for 10 minutes (without HU) or with 100 μM HU for 20 minutes. 50 DNA fiber clusters containing four continuous BrdU-labeled fork replicon in each cluster were analyzed to measure the average intra-cluster fork spacing.

### Neutral comet assay

The neutral comet assay was performed as described ^55^. Glass slides were coated with 0.8% agarose for 5 seconds and air dried. Single cell suspension (∼10^4^ cells/10 μL) were mixed with 70 μL 0.8% low-melting-point agarose (LMP, Sangon Biotech, A600015-0025) kept at 37°C. Cell-agarose suspension was spread onto the prepared slides and covered with coverslip. After being kept at 4°C for 10 minutes, slides were incubated in lysis solution (2.5 M NaCl, 100 mM Na_2_EDTA, 10 mM Tris, 1% N-lauroylsarcosine, 1% TritonX-100, pH =9.5) for 60 minutes at 4°C in the dark without coverslip, then washed and incubated with cold electrophoresis buffer (300 mM sodium acetate, 100 mM Tris, pH =8.3) for 20 minutes, then electrophoresis was performed for 30 minutes at 1 V/cm, 80 mA. Slides were washed with neutralization buffer (0.4 M Tris-HCl, pH 7.4) and fixed in ethanol until air dried. Slides were stained with DAPI (10 ng/mL) and immediately analyzed. Comets were analyzed with Komet 7 comet assay software (Andor Technology). At least 200 cells were analyzed in three independent replications.

### Mitotic chromosomal defects analysis

Cells were cultured under normal conditions on matrigel-coated glass slides (ZEHAO, 12-545-82). When the confluence reached about 70%, cells were fixed with 4% paraformaldehyde for 15 minutes at room temperature and stained with DAPI. 50 mitotic cells were analyzed for each sample to detect chromosome bridge and micronuclei. Each experiment was repeated three times independently.

### Karyotyping

Cells were cultured under normal conditions until about 70% confluence, treated with 120 ng/mL KARYOMAX Colcemid (Gibco, 15212-012) for 2 hours. Cells were then digested with 0.05% trypsin-EDTA (Invitrogen, 25200072), resuspended with hypotonic buffer (10 mM Tris-HCl, pH 7.4; 40 mM glycerol; 20 mM NaCl; 1.0 mM CaCl_2_ and 0.5 mM MgCl_2_) for 15 minutes at 37°C to swell the cells, followed by fixation with methanol/glacial acetic acid (3:1) for 30 minutes. Cells were dropped onto ice-cold wet glass slides, air dried, incubated at 37°C for 24 hours, and stained with 3% Giemsa solution (Gibco, 10092013) at pH 6.8 for 10 minutes. Images were captured using a Leica TCS SP5 confocal microscope system (Leica Microsystems). At least 50 metaphases were analyzed in three independent replications.

### In vivo RNA pull down

The RNA pull down in vivo was performed as described ^52^. DNA probes labeled with biotin at 3’ end were purchased from Guangzhou RiboBio Co.,LTD. In order to perform in vivo RNA pull down, 10^8^ cells were prepared and cross-linked with 265 nm UV light at 400 mJ energy in ice-cold PBS. Then the cells were incubated in CSKT buffer with 1mM PMSF (Beyotime, ST505) and SUPERaseIn (Invitrogen, AM2684) at 4°C for 10 minutes, centrifuged at 1200×g for 10 minutes. The pellet was treated with 3 mL DNase I buffer (50 mM Tris pH 7.5, 0.1% sodium lauroyl sarcosine, 0.5% Nonidet-P 40, 1 × protease inhibitors, 600 U RNase free DNase I, SUPERaseIn, 10 mM vanadyl ribonucleoside complex) at 37°C, centrifuged and discarded the precipitation. The supernatant was pre-cleared using 50 μL M-280 streptavidin Dynabeads (Thermo, 00781251) for 20 minutes at room temperature, and incubated with 100 pmol probes and 160 μL beads at 65°C for 15 minutes. After cool down gradually to 37°C, the beads were washed for three times with wash buffer 1 (50 mM Tris, pH 7.5, 1% SDS, 0.3 M LiCl, 0.5% Nonidet-P 40, 1 mM PMSF, 1 mM DTT, 1 × protease cocktail inhibitors) at 37°C, incubated with 20 U DNase I for 10 minutes at 37°C, washed twice with wash buffer 1 and once with wash buffer 2 (1% SDS, 5 mM EDTA, 1 mM DTT, 150 mM NaCl, 1 mM PMSF). The beads were boiled at 100°C in 2 × SDS protein loading buffer. The proteins were used for mass spectrometry or western blot analysis.

### Immunoprecipitation

10^7^ cells were collected and lysed in RIPA buffer (Beyotime, P0013J) including 1x protease inhibitor (Beyotime, P1006). The lysate was incubated with 40 μL Protein G Dynabeads (Thermo, 88847) and 2 μg antibody at 4°C overnight. After that, the supernatant was removed and the magnetic beads were washed three times. The beads were boiled at 100°C in 2 × SDS protein loading buffer for 10 minutes. The supernatant was used for mass spectrometry analysis or immunoblotting.

### Ibuprofen rescue

*Lnc956^-/-^* males are mated with *Lnc956^+/-^* females. As described ^32^, pregnant females at E7.5 were provided with 250 mL drinking water containing 100 mg ibuprofen (child Advil). Pregnant *Lnc956*^+/-^ female mated with same male was used as control, for which no drug was included in drinking water.

### Statistical analysis

Data were analyzed using GraphPad Prism 7 (GraphPad Software, La Jolla, CA, United States) through two-tailed Student’s *t*-test. P<0.05 was considered significant. All data were provided as the mean ± SEM.

### Data availability

The data of this study are available from the corresponding authors upon reasonable request. RNA-seq data for iROND samples and E12.5 embryos from WT and *Lnc956* KO mice have been deposited in Gene Expression Omnibus database under accession numbers GSE186663 and GSE196639, respectively. The sequence of full-length *Lnc956* transcript can be downloaded from the NONCODE database (http://www.noncode.org/).

## Supporting information

Supplementary infomation

## Acknowledgments

We thank Professor Guohong Li at the Institute of biophysics, CAS for helpful comments, and Yuqi Ning in our lab for deriving *Lnc956* knockout ESC lines. This work was supported by National Natural Science Foundation of China (31930027 to P.Z. and 32000422 to W.D.Z.), National Key Research & Developmental Program of China (2021YFA1102002), Natural Science Foundation of Yunnan province (202001AT070140 to W.D.Z.), and the exchange program of State Key Laboratory of Genetic Resources and Evolution, Kunming Insititute of Zoology, Chinese Academy of Sciences (GREKF21-14 to Z.L.C.).

## Author contributions

Weidao Zhang and Min Tang performed most of the experiments, analyzed the data, and prepared the figures. Lin Wang analyzed the RNA sequencing data and obtained the full-length of *Lnc956.* Hu Zhou and Jing Gao performed the mass spectrometry analysis. Zhongliang Chen constructed some expression plasmids. Bo Zhao developed the iROND method, identified *Lnc956* and supervised the figure preparation. Ping Zheng supervised the study and wrote the manuscript.

## Competing interests

The authors declare no competing interests.

## Notes

### Competing Interest Statement

The authors have declared no competing interest.

