## Supplementary infomation for "*Lnc956*-TRIM28-HSP90B1 complex on replication forks promotes CMG helicase retention to ensure stem cell genomic stability and embryogenesis"

**Supplementary information**

Supplementary information contains:

Supplementary figures 1-7

Supplementary Tables 1-3

Supplementary figures 1-7.

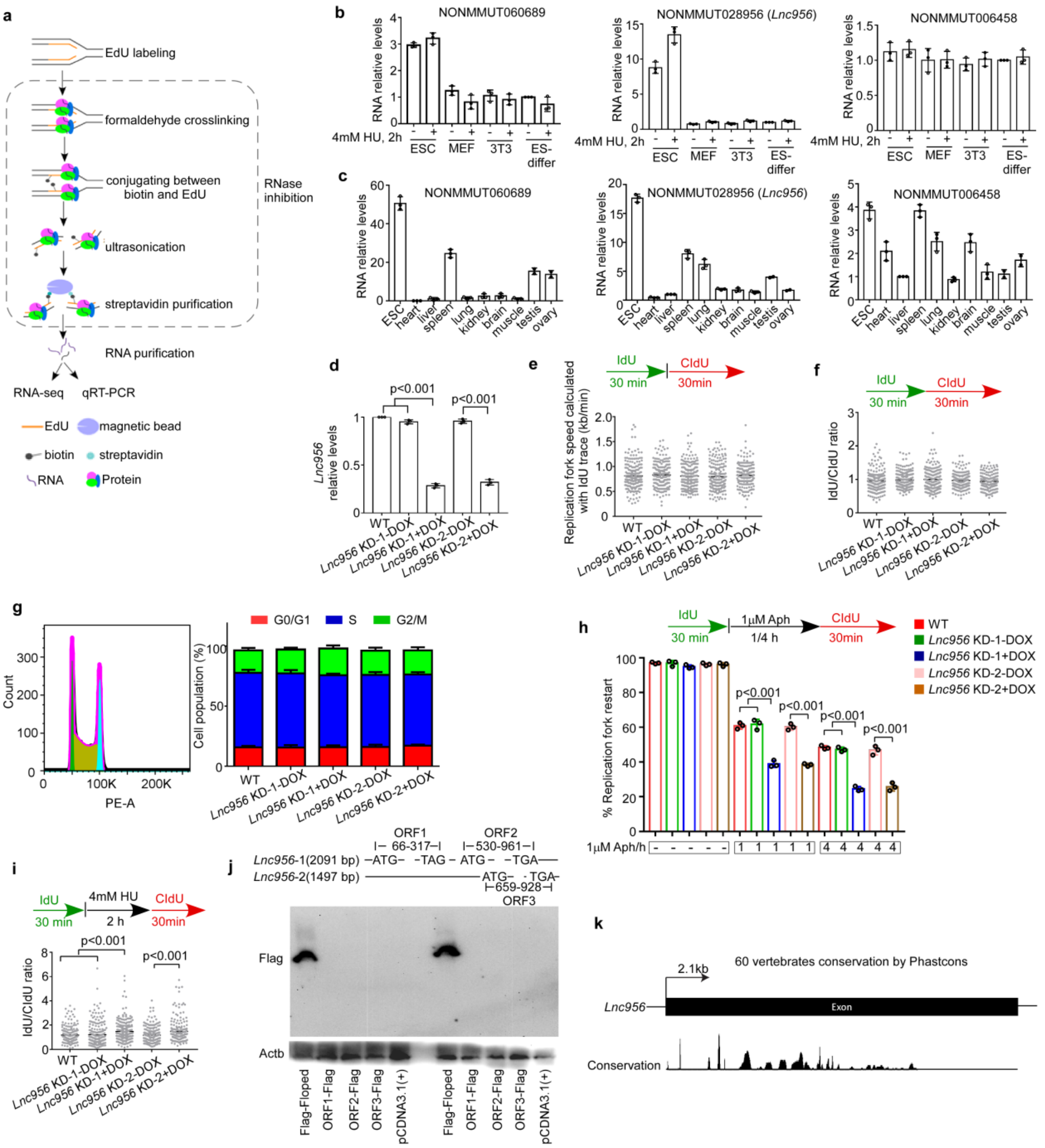


**Supplementary Fig. 1. Identification of ESC-specific lncRNAs localized on replication forks.**

**a** The schematic diagram of iROND (isolate RNA on nascent DNA). **b-c** Quantitative RT-PCR (qRT-PCR) analysis of the relative expression levels of three lncRNAs in mouse ESCs compared to several differentiated cells (**b**) or tissues (**c**). In b, ESCs were differentiated (ES-differ) in the presence of retinoid acid (RA). LncRNA expression was normalized by *Actb*. The relative expression level in ES-differ (b) or in liver (c) was set as 1. **d** qRT-PCR confirmed the efficient knock down (KD) of *Lnc956* in mouse ESCs by two independent DOX-inducible shRNAs. **e-g** Under normal culture condition, *Lnc956* KD did not influence the replication fork speed calculated with IdU trace (e), the fork symmetry reflected by the IdU/CIdU ratio (f), or the cell cycle phase distribution measured by flow cytometry (g). **h** *Lnc956* KD in ESCs compromised the stalled fork restart after treatment with aphidicolin (Aph). **i** *Lnc956* KD in ESCs increased the fork asymmetry under replication stress condition. **j** Three coding fragments in *Lnc956* did not encode protein or peptide. The Flag-tagged Floped protein was used as a positive control. **k** Sequence conservation of *Lnc956* in 60 vertebrates predicted by UCSC Genome Browser track (Phastcons). All data were shown as mean±SEM, two-tailed Student's *t*-test. All experiments were repeated three times with similar results. In (e, f, h, i), at least 200 fibers from three independent experiments were analyzed.

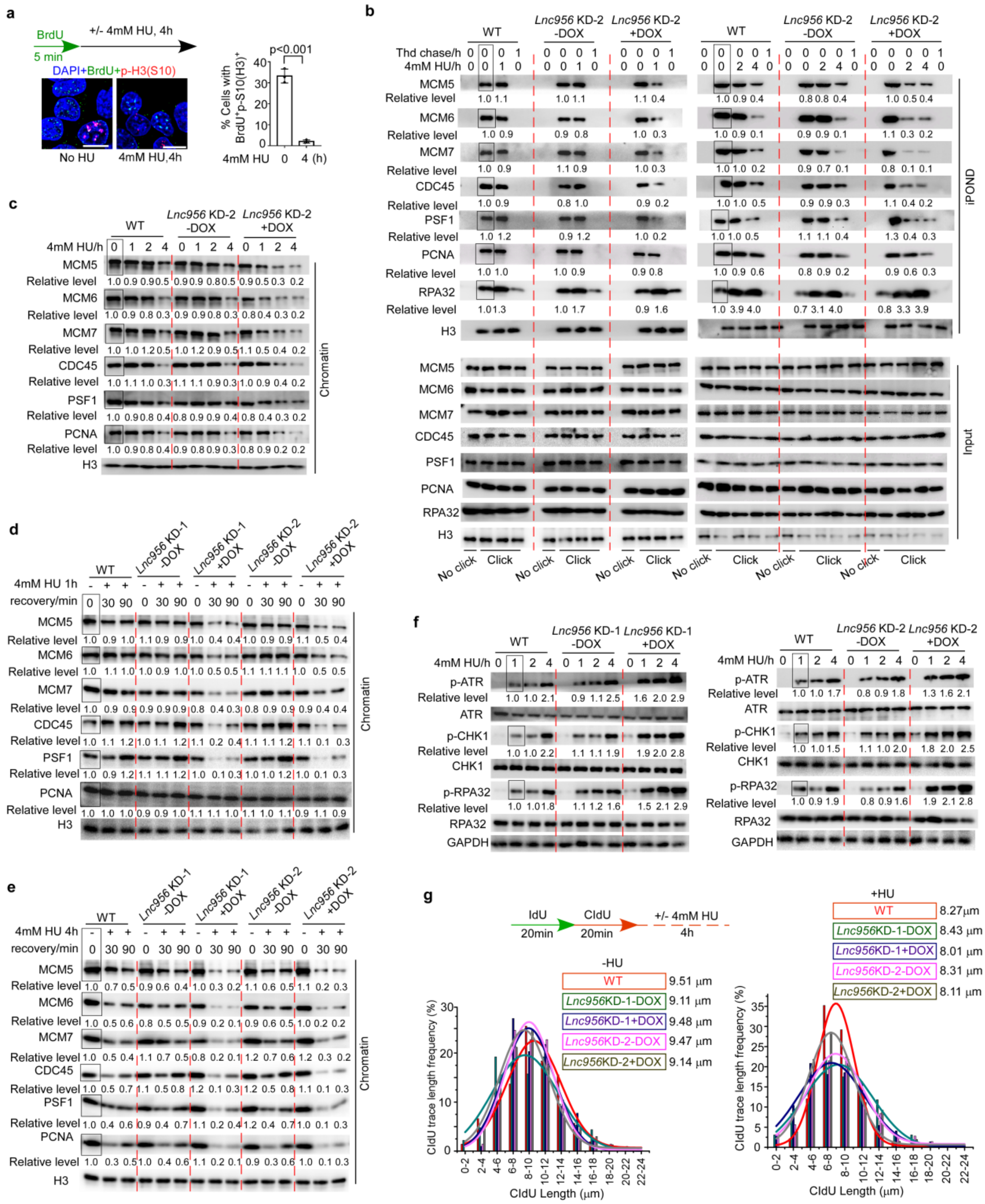


**Supplementary Fig. 2. *Lnc956* KD accelerated the unloading of CMG components without affecting ATR-CHK1 signaling.**

**a** Mouse ESCs were labeled with BrdU for 5 min to mark the S phase cells. The progression of S phase into G2/M phase (monitored by phosphorylation of histone H3 at serine 10 (p-H3(S10)) was measured after 4 h. Treatment with 4 mM HU for 4 h prevented the progression of S phase into G2/M phase.

Scale bar, 10 μm. **b** *Lnc956* knockdown via the second independent shRNA (KD-2) led to premature dissociation of CMG components (MCM5, 6, 7, CDC45 and PSF1) from replication forks under replication stress condition. This defeat was detected at as early as 1 h following HU treatment. Other replication fork proteins PCNA and RPA32 displayed different change patterns. Samples without click reaction (no click) and with 10 μM thymidine (Thd) chase were included as negative controls for iPOND. **c** *Lnc956* KD-2 accelerated the dissociation of CMG components from chromatin. **d-e** ESCs were treated with 4 mM HU for 1 h (**d**) or 4 h (**e**), followed by recovery for 30 min or 90 min. *Lnc956* KD consistently induced the premature dissociation of CMG components from chromatin. **f** ATR-CHK1 signaling was normally activated in *Lnc956* KD ESCs, as indicated by the phosphorylation of ATR at Ser428, CHK1 at Ser345, and RPA32 at Ser33. **g** DNA fiber assay showed that the nascent DNA remained intact in *Lnc956* KD ESCs under normal (left panel) or HU treatment condition (right panel). At least 200 fibers from three independent experiments were analyzed. All experiments were repeated three times with similar results. The relative protein levels were normalized by histone H3 in (b-e) or by GAPDH in (f). The levels in samples marked with box were set as 1.

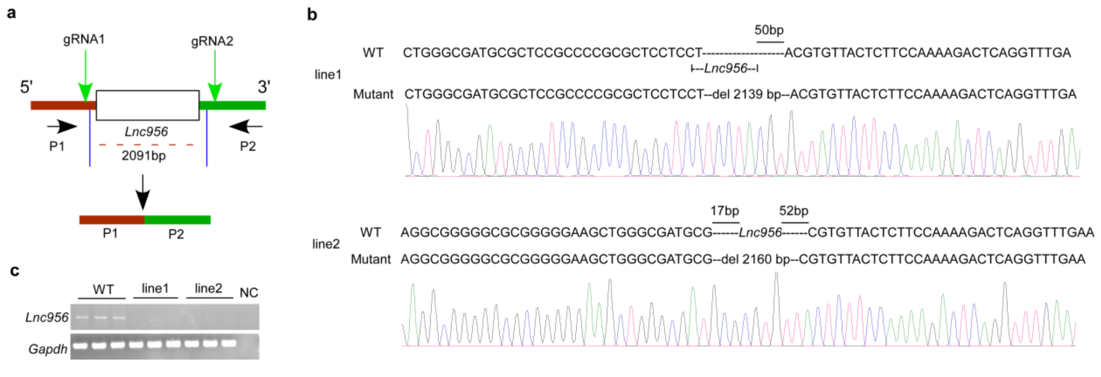


**Supplementary Fig. 3. Generation of *Lnc956* knockout (KO) mice.**

**a** Schematic diagram of CRISPR/Cas9 design to generate *Lnc956* KO mice. P1 and P2 represent primers used for PCR-based genotyping. **b** The genotype of *Lnc956* KO mice was verified by Sanger sequencing of PCR-amplified genomic fragments. Line 1 deleted a 2139-bp fragment, and line 2 contained a 2160-bp fragment deletion. **c** Genotyping of *Lnc956* KO mice by regular PCR.

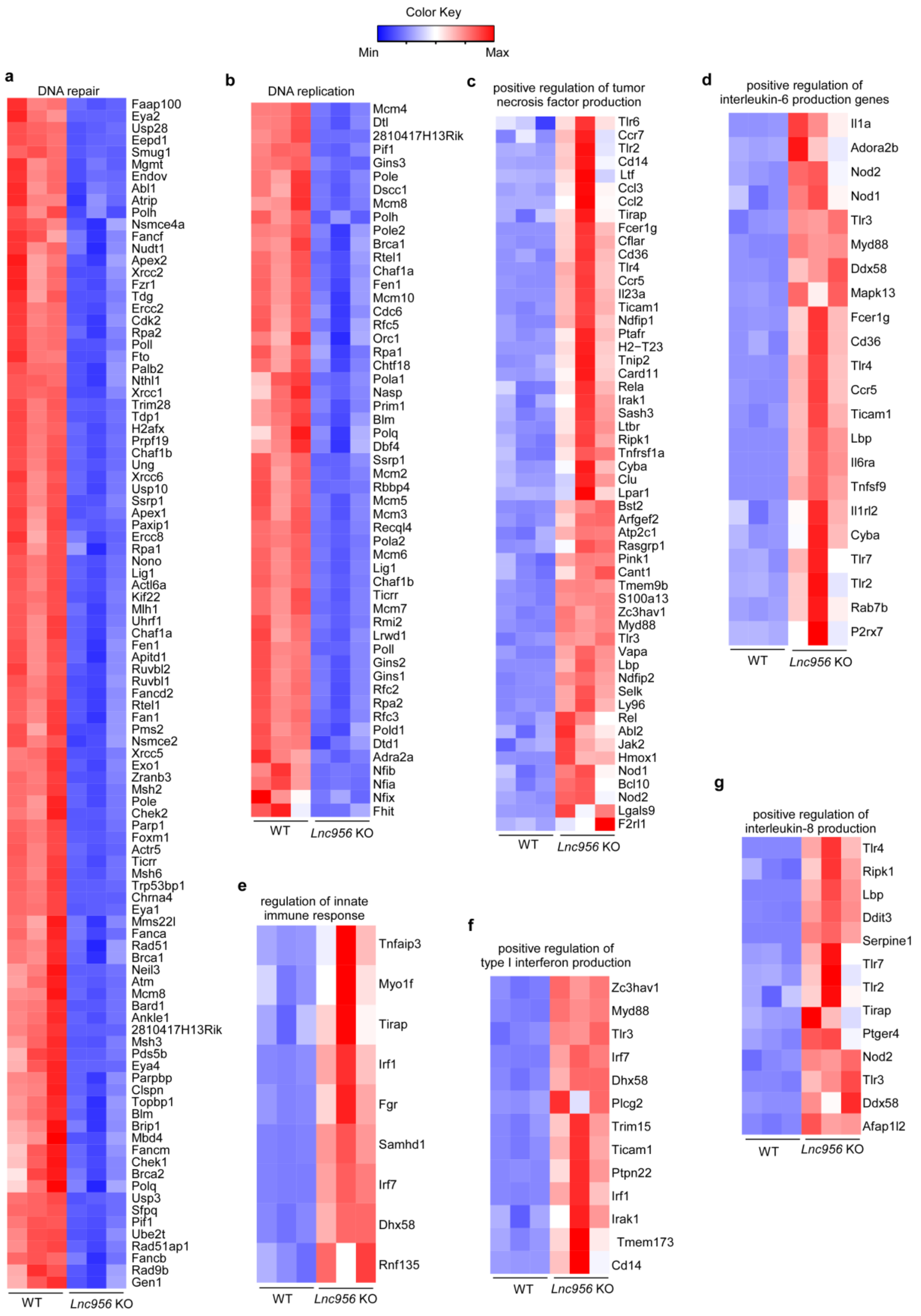


**Supplementary Fig. 4. Gene ontology (GO) enrichments of differentially expressed genes between wildtype (WT) and *Lnc956* knockout (KO) embryos.**

Differentially expressed genes (DEGs) between WT and *Lnc956* KO embryos at E12.5 were analyzed by RNA-seq. Heatmaps of DEGs involved in the GO processes including: (**a**) DNA repair, (**b**) DNA replication, (**c**) positive regulation of tumor necrosis factor production, (**d**) positive regulation of interleukin-6 production, (**e**) regulation of innate immune response, (**f**) positive regulation of type-1 interferon production, and (**g**) positive regulation of interleukin-8 production.

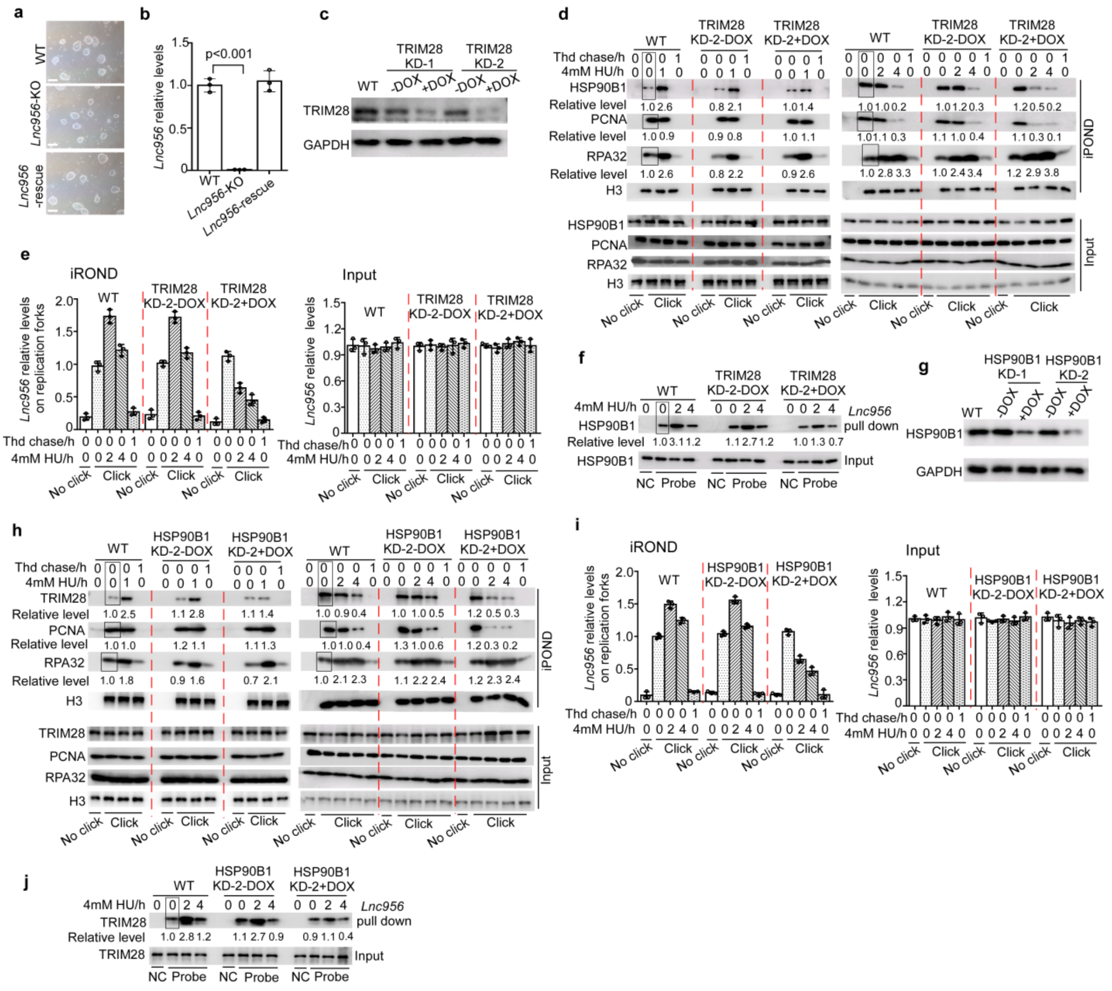


**Supplementary Fig. 5. *Lnc956*-TRIM28-HSP90B1 form a RNP complex in an inter-dependent manner.**

**a** *Lnc956* knockout (KO) ESCs were derived from KO embryos. *Lnc956* was re-expressed in KO ESCs to establish rescue cell line. The KO and rescue ESCs show normal morphology as wildtype (WT) ESCs. **b** qRT-PCR validated the knock out and re-expression of *Lnc956* in KO ESCs. **c** Immunoblotting validated the efficient knockdown (KD) of TRIM28 in ESCs by two independent shRNAs. **d** TRIM28 KD (KD-2) in ESCs reduced the accumulation of HSP90B1 on stalled replication forks upon replication stress. The influence was detected at as early as 1 h after HU treatment. **e** qRT-PCR analysis of iROND samples revealed that TRIM28 KD (KD-2) decreased the accumulation of *Lnc956* on stalled forks. The relative levels of *Lnc956* in inputs were normalized by *Actb*. **f** TRIM28 KD (KD-2) also reduced the association of *Lnc956* with HSP90B1 as revealed by in vivo RNA pulldown. **g** Immunoblotting validated the efficient KD of HSP90B1 in ESCs by two independent shRNAs. **h-j** Likewise, HSP90B1 KD (KD-2) decreased the accumulation of TRIM28 (**h**) and *Lnc956* (**i**) on stalled replication forks, and impaired the TRIM28-*Lnc956* association (**j**) under HU treatment condition. In vivo RNA pulldown using sense probe was set as negative control (NC) in (f, j). The relative protein levels were normalized by histone H3 (d, h), or by input (f, j). The levels in samples marked with box were set as 1. The experiments were repeated three times with consistent results.

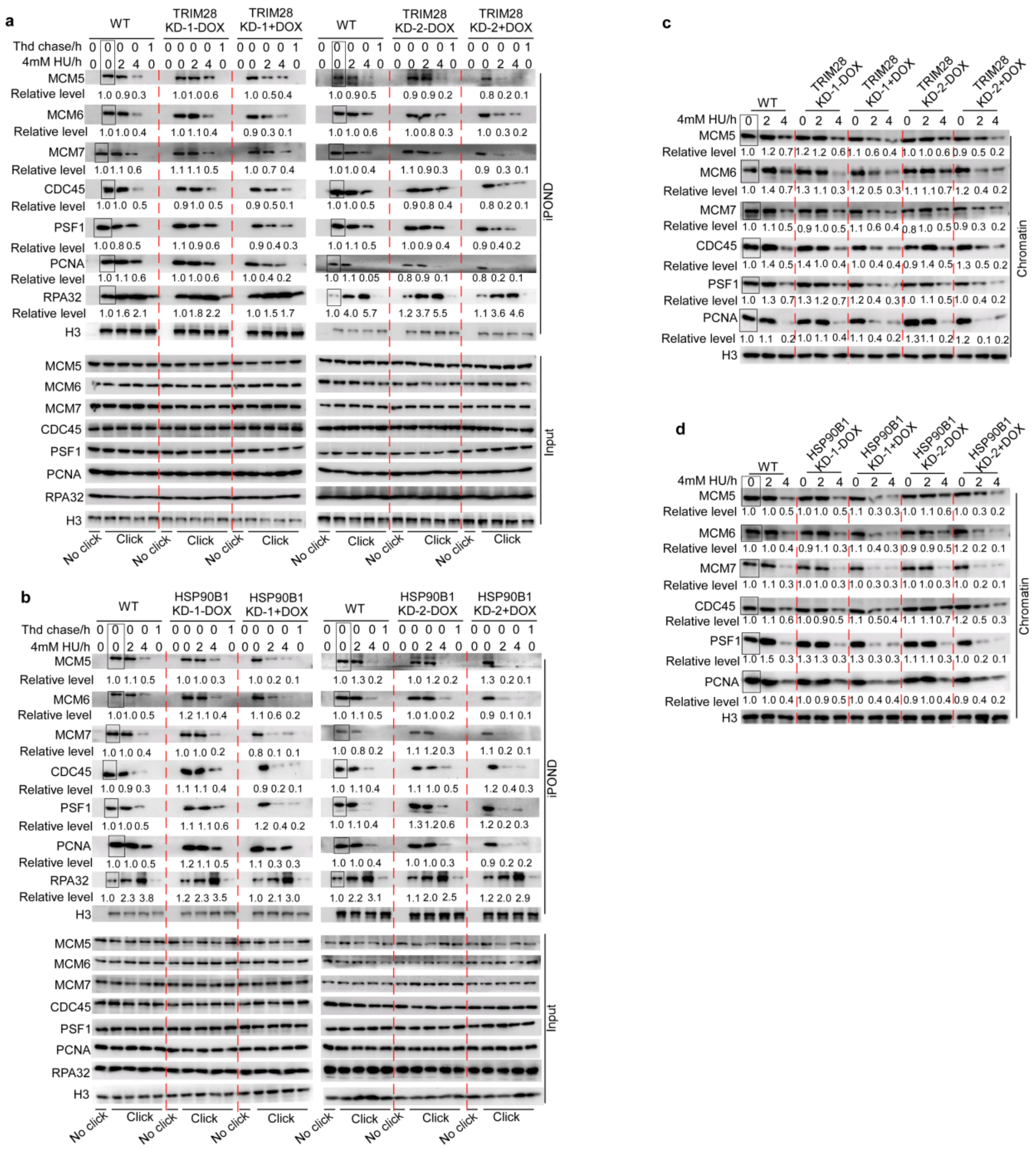


**Supplementary Fig. 6. Knockdown of TRIM28 or HSP90B1 similarly caused the premature unloading of CMG.**

**a-b** iPOND analysis revealed that knockdown (KD) of TRIM28 (**a**) or HSP90B1 (**b**) caused premature unloading of CMG components from stalled replication forks after HU treatment. **c-d** Immunoblotting analyses of purified insoluble chromatin also showed the accelerated dissociation of CMG components from chromatin in TRIM28 KD (**c**) or HSP90B1 KD (**d**) ESCs. The relative protein levels were normalized by histone H3, and the levels in samples marked with box were set as 1. All experiments were repeated three times with similar results.

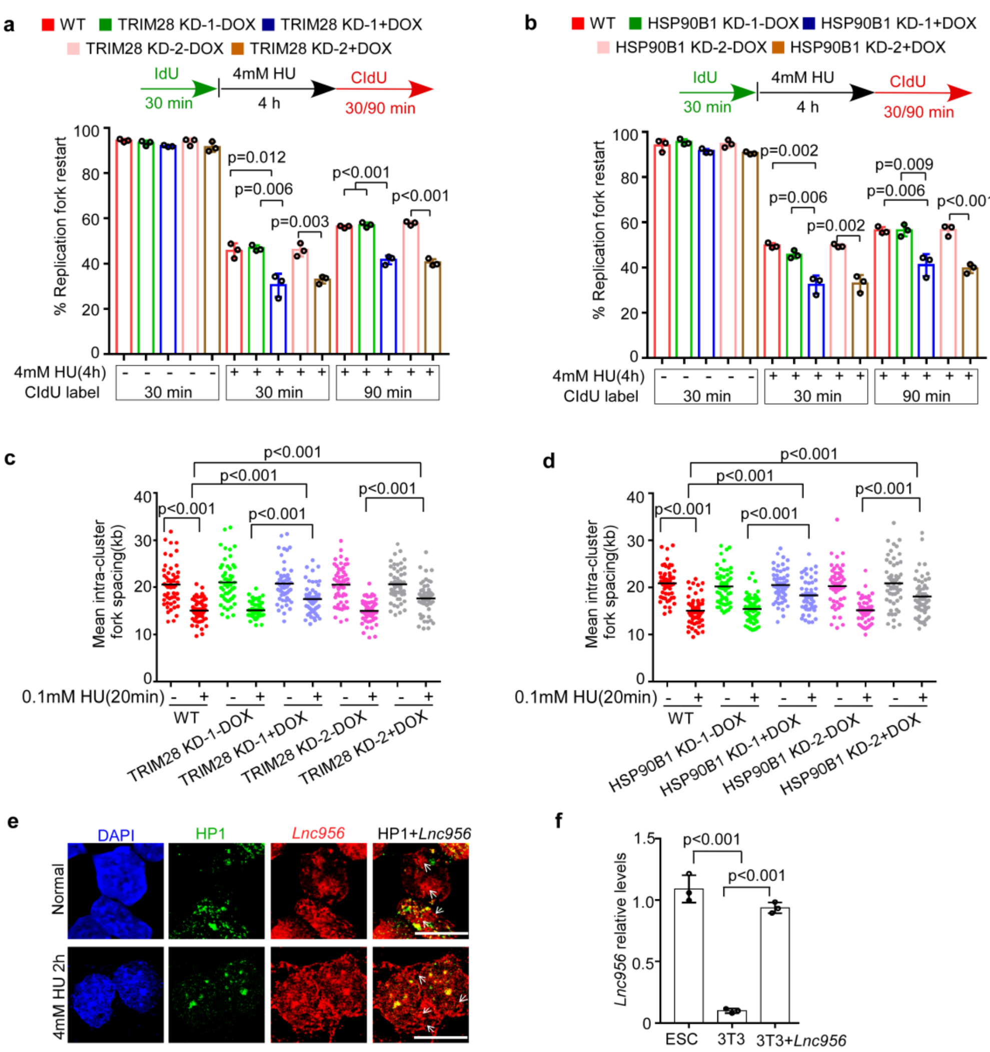

**Supplementary Fig. 7. Knockdown of TRIM28 or HSP90B1 caused replication fork defects.**

**a-b** Two independent KD of TRIM28 (**a**) or HSP90B1 KD (**b**) compromised stalled fork restart. At least 200 fibers from three independent experiments were analyzed. **c-d** KD of TRIM28 (**c**) or HSP90B1 (**d**) increased the inter-origin distances, indicative of reduced dormant origin density. At least 50 continuous fibers from three independent experiments were analyzed. **e** Co-staining of *Lnc956* with HP1 (marker of heterochromatin) showed partial co-localization. Arrows indicated the *Lnc956* foci not co-localized with HP1.

Scale bar, 10 μm. **f** The relative level of *Lnc956* forced expressed in NIH3T3 cells measured by qRT-PCR. *Lnc956* expressions were normalized by *Actb*.

**Supplementary Table 1. Primers for PCR cloning and RT-PCR.**

| Gene | RT-PCR primers | |
|---|---|---|
| *Lnc956* | Forward | 5'-TAGGATCGGCACCTTCTCTGA-3' |
| | Reverse | 5'-CCTTTCACCTTGTGCGACCT-3' |
| NONMMUT060689 | Forward | 5'-GTTTATGCTGCGTCCACTGC-3' |
| | Reverse | 5'-ATAACCGTCAGCTTTGCCCT-3' |
| NONMMUT006458 | Forward | 5'-GGTCTTTCTCGTCCTAGCCG-3' |
| | Reverse | 5'-ATGTGGGTCACCGAAGTGTC-3' |
| *Neat1* | Forward | 5'-TTGGGACAGTGGACGTGTGG-3' |
| | Reverse | 5'-TCAAGTGCCAGCAGACAGCA-3' |
| *Actb* | Forward | 5'-GTGACGTTGACATCCGTAAAGA-3' |
| | Reverse | 5'-GCCGGACTCATCGTACTCC-3' |
| *Gapdh* | Forward | 5'-TTGAGGTCAATGAAGGGGTC-3' |
| | Reverse | 5'-TCGTCCCGTAGACAAAATGG-3' |
| | **RACE primers** | |
| *Lnc956* | 5'-RACE | 5'-GGCAAGCCTAGACCTCTTCAA-3' |
| | 3'-RACE | 5'-CAGCAGCTCAGCGAAGTGGG-3' |
| | **Predicted ORFs vector construction primer** | |
| *Lnc956*-isform1-ORF1 | Forward | 5'-ATGATATCTGGTATCAACGCAGAGTACATG-3' |
| | Reverse | 5'-GCTCTAGACTACTTATCGTCGTCATCCTTGTAATC ATCCCAGGAGTGTGAGGGCT-3' |
| *Lnc956*-isform1-ORF2 | Reverse | 5'-GCTCTAGACTACTTATCGTCGTCATCCTTGTAATC ACCAAGCGTGGGCCACGGGT-3' |
| *Lnc956*-isform2-ORF3 | Reverse | 5'-GCTCTAGACTACTTATCGTCGTCATCCTTGTAATC CACAAACCCAGCAGGTACATAC-3' |
| | **PCR primers** | |
| *Lnc956* KO mouse genotyping | Forward | 5'-CCTCGGAAGACCACCATACTCT-3' |
| | WT/Heter Reverse | 5'-GGACCAGAAGAAGCAGTACGAAG-3' |
| | KO Reverse | 5'-GGCAGCTTATTATTTTTGTTCTTTC-3' |
| *Lnc956* isform | Forward | 5'-TAGGATCGGCACCTTCTCTGA-3' |
| | Reverse | 5'-CCATGTGGTTCCTGGATATT-3' |
| *Sry* | Forward | 5'-ATGTCAAGCGCCCCATGAAT-3' |
| | Reverse | 5'-CCCTCCGATGAGGCTGATATTTA-3' |

**Supplementary Table 2. Short hairpin RNA sequences used for RNA knockdown.**

| **Gene** | **shRNA mir** |
|---|---|
| *Lnc956* KD-1 | 5'-TGCTGTTGACAGTGAGCGATCGCACAAGGTGAAAGGCGAGTAGTGAAGCCACAGATGTACTCGCCTTTCACCTTGTGCGACTGCCTACTGCCTCGGA-3' |
| *Lnc956* KD-2 | 5'-TGCTGTTGACAGTGAGCGAAGTCTCCAGCGCCAAATAAAGTAGTGAAGCCACAGATGTACTTTATTTGGCGCTGGAGACTGTGCCTACTGCCTCGGA-3' |
| *Hsp90b1* KD-1 | 5'-TGCTGTTGACAGTGAGCGAATTATACTCTCGCTATGAATCTAGTGAAGCCACAGATGTAGATTCATAGCGAGAGTATAATGTGCCTACTGCCTCGGA-3' |
| *Hsp90b1* KD-2 | 5'-TGCTGTTGACAGTGAGCGAAGGAGCTTAGACTCGGGATTGTAGTGAAGCCACAGATGTACAATCCCGAGTCTAAGCTCCTGTGCCTACTGCCTCGGA-3' |
| *Trim28* KD-1 | 5'-TGCTGTTGACAGTGAGCGAAGACAGAAGCTGCTATTGGAGTAGTGAAGCCACAGATGTACTCCAATAGCAGCTTCTGTCTCTGCCTACTGCCTCGGA-3' |
| *Trim28* KD-2 | 5'-TGCTGTTGACAGTGAGCGCTTACTTCTGTGGATTTAATAATAGTGAAGCCACAGATGTATTATTAAATCCACAGAAGTAAATGCCTACTGCCTCGGA-3' |

**Supplementary Table 3. Antibody information.**

| Antibodies | Source | Dilution |
|---|---|---|
| Mouse monoclonal anti-BrdU (clone B44), also used for immunostaining of IdU | BD Biosciences; Cat#347580 | 1:500 (IF) |
| Rat monoclonal anti-BrdU (clone BU1/75), also used for immunostaining of CldU | Novus; Cat#NB500-169 | 1:1000 (IF) |
| Mouse monoclonal anti-PCNA (clone PC10) | Cell Signaling Technology; Cat#2586 | 1:1000 (IB) |
| Ds DNA maker (HYB331-01) | Santa Cruz; Cat#sc-58749 | 1:1000 (IB) |
| Mouse monoclonal anti-GAPDH | Sangon Biotech; Cat#D190090 | 1:1000 (IB) |
| Rabbit polyclonal anti-γ-H2AX (clone 20E3) | Cell Signaling Technology; Cat#9718 | 1:1000 (IB) |
| CHK1 Mouse monoclonal antibody (clone 2G1D5) | Cell Signaling Technology; Cat#2360S | 1:1000 (IB) |
| Rabbit anti-p-ATR (Ser428) | Cell Signaling Technology; Cat#2853S | 1:1000 (IB) |
| Rabbit polyclonal anti-ATR | Cell Signaling Technology; Cat#2790S | 1:1000 (IB) |
| P-CHK1(S345) Rabbit monoclonal antibody(133D3) | Cell Signaling Technology; Cat#2348S | 1:1000 (IB) |
| Anti-RPA32/RPA2 antibody | Abcam; Cat#ab76420 | 1:1000(IB) |
| Anti-RPA32/RPA2 (phosphoS33) antibody | Abcam; Cat#ab211877 | 1:1000(IB) |
| β-Actin mouse antibody | Abclonal; Cat#AC004 | 1:10000 (IB) |
| Histone H3 (D1H2) XP® Rabbit mAb | Cell Signaling Technology; Cat#4499S | 1:1000(IB) |
| Anti-KAP1(TRIM28) antibody | Abcam; Cat#ab10483 | 1:1000 (IB) |
| Recombinant Anti-MCM5 antibody (clone EP2683Y) | Abcam; Cat#ab75975 | 1:1000 (IB) |
| Recombinant Anti-MCM6 antibody (clone EPR17686) | Abcam; Cat#ab201683 | 1:1000 (IB) |
| MCM7 XP Rabbit mAb (clone D10A11) | Cell Signaling Technology; Cat#3735T | 1:1000 (IB) |
| Recombinant Anti-HSP90B1 (GRP94) antibody | Abcam; Cat#ab238126 | 1:1000 (IB) |
| CDC45 Rabbit mAb (clone D7G6) | Cell Signaling Technology; Cat#11881s | 1:1000 (IB) |
| Recombinant Anti-PSF1 antibody (clone EPR13359) | Abcam; ab181112 | 1:1000 (IB) |
| Rabbit monoclonal anti-K63-linkage Specific Polyubiquitin (clone D7A11) | Cell Signaling Technology; Cat#5621 | 1:1000 (IB) |
| K48-linkage Specific Polyubiquitin (cloneD9D5) Rabbit mAb | Cell Signaling Technology; Cat#8081 | 1:1000 (IB) |
| HP1β XP® Rabbit mAb(cloneD2F2) | Cell Signaling Technology; Cat#8676 | 1:500 (IF) |
| Phospho-Histone H3 (Ser10) XP® Rabbit mAb (cloneD7N8E) | Cell Signaling Technology; Cat#53348 | 1:1000 (IF) |
| Goat anti-Mouse IgG (H+L) Secondary Antibody, Alexa Fluor 488 | Thermo Fisher Scientific; Cat#A-11029 | 1:500 (IF) |
| Goat anti-Rat IgG Secondary Antibody, Alexa Fluor Cy3 | Thermo Fisher Scientific; Cat#A-10522 | 1:500 (IF) |
| Goat anti-Rat IgG (H+L) Secondary | Thermo Fisher Scientific; Cat#A-21247 | 1:500 (IF) |

| | | |
|---|---|---|
| Antibody, Alexa Fluor 647 | | |
| Goat anti-Rabbit IgG (H+L) Secondary Antibody, Alexa Fluor 488 | Thermo Fisher Scientific; Cat#A-32731 | 1:500 (IF) |
| Goat anti-Mouse IgG (H+L) Secondary Antibody, HRP | Thermo Fisher Scientific; Cat#31430 | 1:5000 (IB) |
| Goat anti-Rabbit IgG (H+L) Secondary Antibody, HRP | Thermo Fisher Scientific; Cat#31460 | 1:5000 (IB) |
